# Ensemble remodeling supports memory-updating

**DOI:** 10.1101/2022.06.02.494530

**Authors:** William Mau, Austin M. Baggetta, Zhe Dong, Brian M. Sweis, Denisse Morales-Rodriguez, Zachary T. Pennington, Taylor Francisco, David J. Freedman, Mark G. Baxter, Tristan Shuman, Denise J. Cai

## Abstract

Memory-updating is critical in dynamic environments because updating memories with new information promotes versatility. However, little is known about how memories are updated with new information. To study how neuronal ensembles might support memory-updating, we used a hippocampus-dependent spatial reversal task to measure hippocampal ensemble dynamics when mice switched navigational goals. Using Miniscope calcium imaging, we identified neuronal ensembles (co-active neurons) in dorsal CA1 that were spatially tuned and stable across training sessions. When reward locations were moved during a reversal session, a subset of these ensembles decreased their activation strength, correlating with memory-updating. These “remodeling” ensembles were a result of weakly-connected neurons becoming less co-active with their peers. Middle-aged mice were impaired in reversal learning, and the prevalence of their remodeling ensembles correlated with their memory-updating performance. Therefore, we have identified a mechanism where the hippocampus breaks down ensembles to support memory-updating.

## Introduction

Memory-updating is important for behavioral adaptations to dynamic environments. Memory recall can use past experiences to guide future behavior, but the ability to forget the past and integrate new information is also critical for adapting to changing circumstances, especially if past information is unreliable or no longer valid^1–4^. The importance of this ability is highlighted by mnemonic and behavioral inflexibility seen in many neurological and psychiatric conditions^5–14^.

Therefore, it is critical to understand the neural underpinnings of memory-updating, not only to address these pathologies, but also to gain a holistic understanding of memory function in general. There is a wealth of knowledge on the neural mechanisms of memory storage, but in comparison, the mechanisms for memory-updating in the brain are poorly understood. Past research has shown that memories are stored in stable “engrams” or neuronal ensembles, groups of neurons that are both necessary and sufficient for memory retrieval^15–19^. However, these ensembles undergo neuroplasticity – remodeling of these ensembles is thought to be the putative mechanism for modifying memories^1–4^, which can support higher-order learning and the ability to flexibly adapt to dynamic environments. In this study, we used a spatial reversal learning task to investigate how hippocampal ensemble dynamics support memory-updating.

Reversal learning is a common paradigm for measuring memory flexibility^20^ where the subject must inhibit previously learned behavioral responses and adopt new ones. Although many reversal learning tasks focus on striatal, orbitofrontal, and prefrontal cortical contributions^20–22^, the hippocampus has also been shown to be important for memory flexibility during spatial reversal^6,23,24^. For example, long-term depression (LTD) in the hippocampus is critical for reversal learning of a spatial memory on the Morris water maze^25–27^. At the circuit level though, it remains unknown how neuronal populations are modified during memory-updating. To answer this question, we hypothesized that during reversal learning, previously stable ensemble activity might break down to support memory-updating of new goal locations and forgetting of old goal locations.

## Results

### Dorsal CA1 neurons do not exhibit global nor rate remapping during Reversal

To examine the real-time dynamics of hippocampal ensembles during memory-updating, we used the latest iteration of miniature head-mounted microscopes (UCLA V4 Miniscopes) to conduct one-photon calcium imaging in dorsal CA1^28–31^ while mice performed a hippocampus-dependent spatial reversal task (Fig. 1a). Prior to behavioral testing, mice received a unilateral (right hemisphere) infusion of AAV1-syn-GCaMP6f-WPRE-SV40 and were implanted with a 1 mm diameter GRIN lens and base plate prior to the experiment. Mice were water-deprived and trained on a circular track with distinct spatial cues to retrieve water rewards from two water ports for four consecutive days (Training1-4). Miniscope mice learned the task across Training1-4, as measured by d’, hit rate, and correct rejection rate as they passed reward sites on each trial, defined as one lap around the circular track (Fig. 1b-d). On the fifth day, we ran a Reversal session, where the two previous ports became unrewarded and two new ports were rewarded. During Reversal, mice licked at incorrect ports more than during Training4, but gradually learned the new goal locations (Supplementary Fig. 1a-d). In subsequent analyses, we chose to focus on correct rejection rate which is an important metric for assessing whether a subject has updated their memory for a reward site being no longer rewarded^32^. Correct rejection rate in particular decreased during Reversal, but hit rate did not (Fig. 1e-f, Supplementary Fig. 1c). During Reversal, mice licked more at previously rewarded ports compared to never-rewarded ports (Fig. 1g) but slowly learned to avoid these ports over the course of the session (Supplementary Fig. 1d). To ensure that hippocampus was necessary for memory-updating during Reversal, we also used the chemogenetic PSAM system^33^ to inactivate the dorsal hippocampus during the Reversal session, which suppressed performance (Supplementary Fig. 2).Thus, with the mice showing memory-updating during the Reversal session, we next examined hippocampal activity.

**Figure 1.**
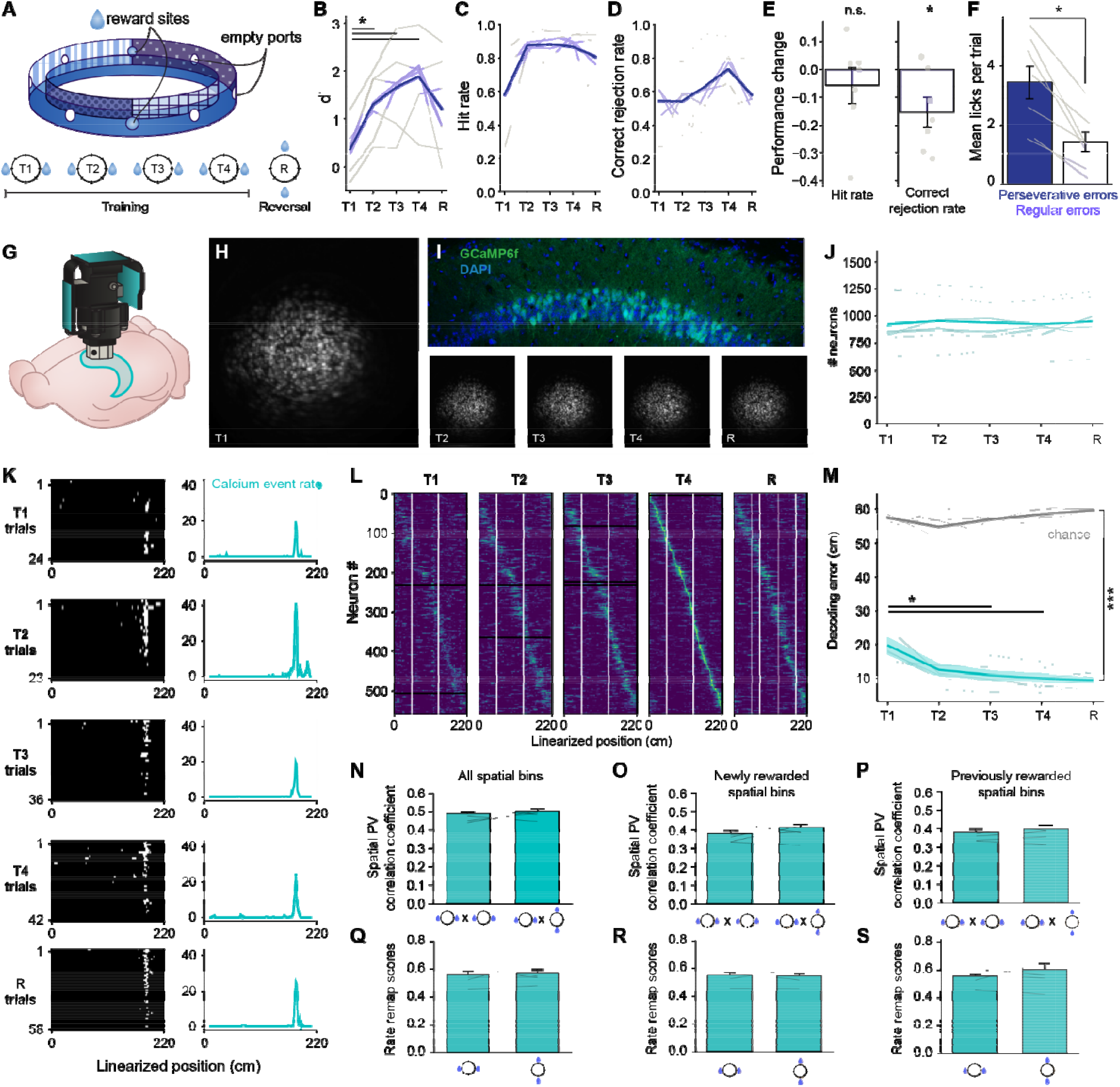
No global or rate remapping during spatial reversal learning. A) Schematic and schedule of the spatial reversal task. Training1-4 are abbreviated as T1-4 throughout. Reversal is abbreviated as R. B) Behavioral d’ of mice across Training and Reversal. Mice improved their performance over sessions (repeated-measures ANOVA F_4,24_=9.17, p=0.0012, *pairwise t-tests p<0.05 after Benjamini-Hochberg corrections for multiple comparisons). C) Hit rate over sessions. Mice improved over sessions (repeated-measures ANOVA F_4,24_=4.77, p=0.0056). D) Correct rejection rate over sessions. Mice improved over sessions (repeated-measures ANOVA F_4,24_=3.15, p=0.032). E) Change in hit rates and correct rejection rates on Reversal compared to Training4 for each mouse. Hit rates did not decrease on Reversal (one-sample t-test compared to 0, t_6_=-0.90, p=0.41). Correct rejection rates decreased relative to Training4 (t_6_=-2.88, p=0.028). F) Mean licks per trial from each mouse during Reversal on previously rewarded ports (perseverative errors, dark blue bar) or never rewarded ports (regular errors, light blue bar). Mice made more perseverative errors compared to regular errors (Wilcoxon signed-ranked test W=0.0, p=0.016). G) Schematic showing in vivo calcium imaging with Miniscopes above dorsal CA1. H) Maximum projections of the field of view in one mouse across sessions. I) Histology of GCaMP6f-expressing cells overlayed with DAPI staining in dorsal CA1. J) Number of recorded neurons on each session per animal. There were no differences in the number of neurons across sessions (repeated-measures ANOVA F_4,24_=0.63, p=0.64). K) Example neuron recorded over all 5 days. Left, spatial activity over trials. Right, trial-averaged tuning curve for that day. L) Trial-averaged activity of the same neurons recorded across Training and Reversal. Neurons are sorted according to their field location on Training4. Only neurons that were registered across all five sessions are plotted. Black rows correspond to neurons that had no activity during periods running above the speed threshold of 7 cm s_-1_. White vertical lines correspond to the rewarded port location. On the Reversal session, the transparent white line corresponds to the reward ports on previous Training days. M) Within-session spatial decoding accuracy using classifiers trained on neuronal activity. All sessions performed better than chance (gray line, permutation tests p<0.001), and there was an improvement in decoding over sessions (repeated-measures ANOVA F_4,24_=16.5, ***p<0.001). Decoding error decreased significantly after Training1 (*pairwise t-tests p<0.04 after Sidak corrections for multiple comparisons). N) Mean spatial population vector (PV) correlation coefficients for each mouse comparing two Training sessions (Training3 x Training4) or a Training session and Reversal (Training4 x Reversal). There was no difference in PV correlations between the two session pairs (Wilcoxon signed-rank test W=8.0, p=0.38). O) Mean spatial PV correlation coefficients across sessions, but restricted to spatial bins 10 cm around the rewarded spatial bins on Training4 and the newly rewarded spatial bins on Reversal. There was no difference between the two session pairs (W=8.0, p=0.38). P) Mean spatial PV correlation coefficients across sessions, but restricted to spatial bins 10 cm around the original set of rewards on both Training4 and Reversal. There was no difference between the two session pairs (W=6.0, p=0.22). Q) Mean rate remapping scores for each mouse on Training4 and Reversal. There was no difference across the two sessions (W=13.0, p=0.94). R) Mean rate remapping scores, but restricted to spatial bins 10 cm around the rewarded ports on the plotted session (original reward bins in Training4 and newly rewarded bins in Reversal). There was no difference between the two sessions (W=10.0, p=0.58). S) Mean rate remapping scores, but restricted to spatial bins 10 cm around the original set of rewards for both Training4 and Reversal. There was no difference between the two sessions (W=8.0, p=0.38).

During behavior, we simultaneously recorded 747.0 ± 42.4 dorsal CA1 neurons per animal, per session (Fig. 1g-j). Dorsal CA1 neurons are well-known for expressing spatial receptive fields (“place fields”)^34^, and consistent with prior studies, we found 39.2 ± 2.4% of the neuronal population fit the criteria for place cells (e.g., Fig 1k; see Methods). The entire neuronal population (Fig. 1l, Supplementary Fig. 3) was relatively stable across Training1-4 and Reversal, reliably predicted spatial location (Fig. 1m), and over-represented reward sites (Supplementary Fig. 1e-g). Based on prior studies where reward locations were suddenly changed^35,36^, we expected hippocampal neurons to change their place field locations (“global remapping”)^37^ or activity rates (“rate remapping”)^38^ during Reversal. However, we did not see more remapping across Training4 and Reversal compared to across Training3 and Training4 (Fig. 1n). This was true even when we restricted our analysis to fields near the new reward during Reversal (Fig. 1o) or the old rewarded sites during Training4 (Fig. 1p). Similarly, activity rates did not change over the course of Reversal, as measured by a rate remapping score (see Methods; Fig. 1q-s). Though we were surprised to not observe widespread population changes during Reversal, this is consistent with other studies showing little to no remapping even during a context change^39,40^. In our task, the spatial cues did not change, which may have contributed to overall spatial stability of place cell tuning curves. Moreover, the traditional approaches for measuring population-wide neural dynamics involves averaging across trials, potentially masking more subtle changes across time. Though there was no evidence of global nor rate remapping during Reversal, it remained possible that the hippocampus was supporting memory-updating by altering the temporal coordination between neurons^39^. Thus, we turned our attention to ensemble dynamics over time during the Reversal session. We chose to agnostically define our ensembles based solely on neuronal co-activity (irrespective of the animal’s corresponding behavior), so we included not only place cells but all recorded neurons to characterize ensemble dynamics in a non-biased manner.

### Dorsal CA1 neurons grouped into neuronal ensembles that showed spatial and reward port selectivity

Past studies have shown that memories are stored in groups of neurons or neural ensembles with their coordinated activity^41,42^. To ask how neuronal ensembles supported spatial memory during Reversal, we first grouped neurons that were co-active into ensembles and then verified that the ensembles operated as expected based on the spatial processing role of the hippocampus. We used an independent component analysis (ICA)-based method^43^ that identified 54.7 ± 3.9 neuronal ensembles per session, each corresponding to groups of neurons that were statistically more co-active than expected by chance. For each ensemble, we computed its weight vector (see Methods), which describes how much each neuron contributes to that ensemble (Fig. 2a), as well as its activation strength (Fig. 2b-c). The number of ensembles did not change over Training1-4 and Reversal (Fig. 2d). Ensembles were sparse; each ensemble was comprised of 3.45 ± 0.06% of the total number of recorded neurons for that session (see Methods for how we defined ensemble membership). The sizes of these ensembles did not change over time (Supplementary Fig. 4a). To verify that these hippocampal ensembles encoded relevant information, we next investigated their spatial selectivity. Using similar criteria that we applied to single neurons, we found that 59.6 ± 1.8% of ensembles showed statistically significant spatial activity (e.g., Fig. 2c,f), in line with other studies showing that neurons in the hippocampus collectively encode spatial information^44,45^. The average spatial information of ensembles increased over Training1-4 (Supplementary Fig. 4b) and we could decode the spatial location of the mice significantly better than chance (Fig. 2e). Collectively, these results demonstrate that we can identify ensembles comprised of sparse, co-firing neuronal populations that contain spatial information, validating our ensemble identification methods.

**Figure 2.**
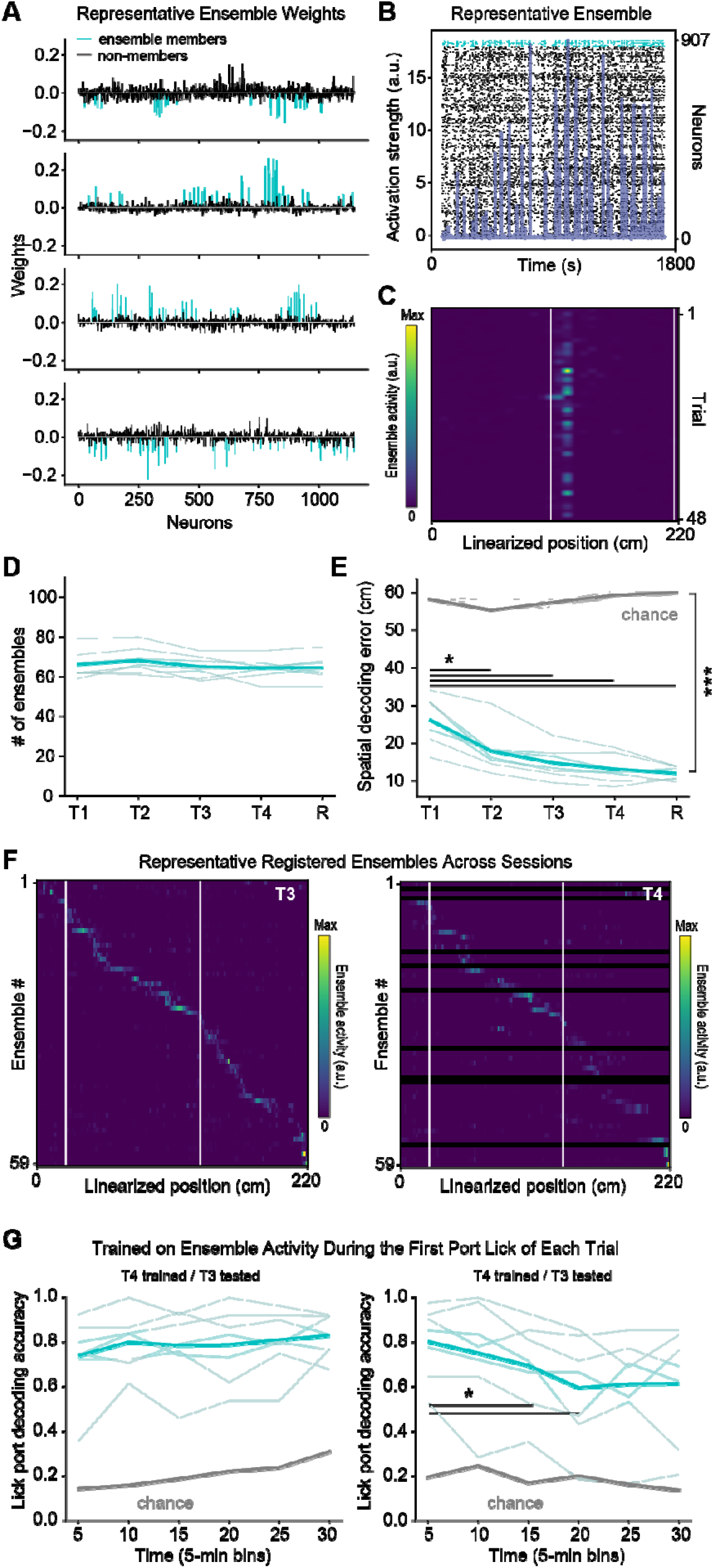
Co-firing CA1 neurons form ensembles and contain spatial and lick information. A) Example weight vectors for four ensembles recorded during Training4. Weights were colored cyan for ensemble members (neurons with weight magnitudes > 2 standard deviations above the mean for that ensemble) or black for non-members. B) Example neuronal ensemble detected during Reversal. The darker blue trace indicates the strength of that ensemble’s activation over time. The raster in the background is sorted by descending neuronal weights. Cyan ticks correspond to calcium transients from ensemble members whereas black ticks are calcium transients from non-members. C) Example of ensemble activity from the same ensemble in B), over trials within a session. D) Number of identified ensembles per animal over sessions. There was no change in ensemble count across sessions (repeated-measures ANOVA F_4,52_=1.60, p=0.19). E) Within-session spatial decoding accuracy using classifiers trained on ensemble activity. All sessions performed better than chance (gray line, permutation tests p<0.001), and there was an improvement in decoding over sessions (repeated-measures ANOVA F_4,24_=29.3, p<0.001). Decoding error decreased significantly after Training1 (pairwise t-tests all p<0.03 after Sidak corrections for multiple comparisons). Chance (gray line) was computed by training classifiers on shuffled data. F) Trial-averaged activity of ensembles registered across Training3 and Training4, ordered by field location on Training3. White lines indicate rewarded port locations. Ensembles on Training3 that had no corresponding match on Training4 appear as black rows in the second plot. G) Lick decoding accuracy using ensembles from Training4 registered to either Training3 (left) or Reversal (right). Classifiers were trained on Training4 ensemble activity during the first lick of each reward port on each trial. There was no significant main effect of session (repeated-measures ANOVA F_1,6_=3.01, p=0.13) or time (F_5,30_=1.76, p=0.15), but a significant interaction (F_5,30_=7.63, p=0.0001). Post-hoc pairwise t-tests revealed significant decreases in decoding accuracy over time only on the Reversal session (*p<0.05 after Sidak’s correction for multiple comparisons). Chance (gray line) was computed by training classifiers on shuffled data.

In addition to spatial location, these ensembles also encoded lick port identity across sessions. In the next analysis, we focused on Training3-4, when the goals were well-learned, and the Reversal session, when goals were switched, to examine whether the ensemble code could predict lick port identities before and after the goals had switched. We registered ensembles across session pairs by matching the ensembles with the highest cosine similarities between their weight vectors (see Methods; Supplementary Fig. 5a-c). We then trained a classifier to decode lick port identity based on ensemble activity collected during the initial lick of each trial at each port. Only the ensemble activity during the initial lick was used to prevent oversampling during reward consumption. When trained on Training4 data and tested on Training3 ensemble activity, the classifier successfully decoded lick port identity as shown by the consistently high decoding accuracy (Fig. 2g, left). When the same classifier was tested on the Reversal session, it was accurate at the beginning of the session, but accuracy decreased over time, suggesting that the old ensemble code for lick port identity was being remodeled (Fig. 2g, right). We found similar results when we instead trained on the ensemble activity 1 s prior to the first lick of each reward port per trial, suggesting that ensemble activity might encode anticipatory actions (Supplementary Fig. 5d). Having observed changes in the ensemble code during Reversal, we next sought to investigate how ensembles collectively remodeled to support memory-updating.

### A subset of dorsal CA1 ensembles remodel during memory-updating and is correlated with enhanced behavioral performance

Based on the reduction in decoding accuracy during Reversal, we hypothesized that ensembles during the Reversal session were undergoing plasticity and remodeling as the mice learned to both lick at the new reward sites and inhibit licking at the old reward sites. Accordingly, we identified a subset of ensembles that decreased their activation strength over the course of the Reversal session. These ensembles whose peak activation strength decreased over trials, we termed “remodeling” ensembles (Fig. 3a). Comparatively, other ensembles during Reversal showed constant activation strength over trials (“non-remodeling” ensembles, Fig. 3b). Because memory-updating occurred during the Reversal session, we expected a higher occurrence rate of remodeling ensembles during Reversal compared to during expression of a stabilized memory at Training4. Indeed, a higher proportion of ensembles were statistically classified as remodeling ensembles during Reversal compared to Training4 (Fig. 3c). This effect was not due to a higher proportion of remodeling neurons (neurons that changed their calcium transient rate over time; Supplementary Fig. 6a), nor a change in the number of active neurons during Reversal (Fig. 1J). Critically, the proportion of remodeling ensembles correlated with the peak behavioral performance (i.e., correct rejection rate) of the mouse during Reversal, suggesting that this ensemble remodeling supports memory-updating (Fig. 3d). Moreover, the rate of ensemble remodeling correlated with the rate of learning to reject old reward locations (Fig. 3e), suggesting a relationship between how quickly the mice learned and how well they were able to adapt their neural representations. The locations where these remodeling ensembles occurred were not more concentrated at any particular reward ports than expected from chance (Supplementary Fig. 6b), refuting the possibility that remodeling ensembles reduced their activity due to decreased visits to the previously-rewarded ports. This suggests that ensemble remodeling is not occurring specifically at specific reward ports, and that it may instead result from circuit-wide remodeling to accommodate incoming conflicting information^46^.

**Figure 3.**
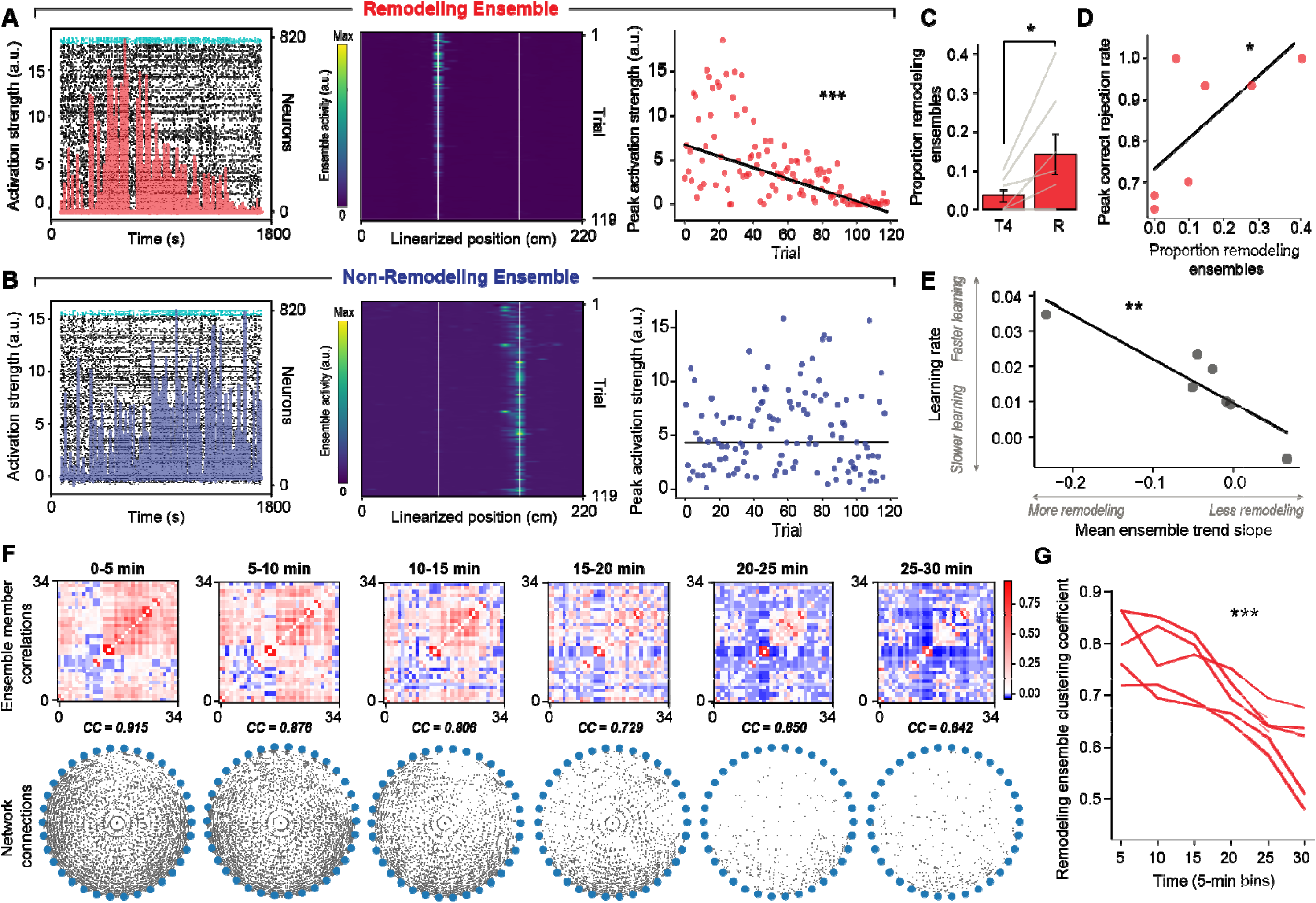
Hippocampal ensembles had reduced stability during Reversal; remodeling ensemble strength correlated with behavioral performance. A) One example “remodeling” ensemble. Left, raster of the remodeling ensemble plotted like Fig. 2B. Center, spatially binned activation strength of the ensemble per trial. White lines indicate rewarded port locations. Right, peak activation strength of the ensemble per trial. ***p<0.001 Mann-Kendall trend test after Sidak’s correction for multiple tests. B) Same as A), but for a non-remodeling ensemble. C) Proportions of remodeling ensembles (out of all detected ensembles) for Training4 (T4) compared to Reversal (R). The proportion of remodeling ensembles increased during Reversal (Wilcoxon signed-rank test, W: 1.5, p=0.034). D) Scatter plot of remodeling ensemble proportion against peak correct rejection rate during Reversal. The proportion of remodeling ensembles was correlated with peak correct rejection rate (Spearman R=0.697, p=0.041). E) Mean ensemble trend slopes from Mann-Kendall trend tests for each mouse against their learning rate (regression slope of the correct rejection rate over trials). The average rate of remodeling correlated with learning rate (Spearman R=-0.89, p=0.0068). F) Within-ensemble cell correlations of one example remodeling ensemble during Reversal. Top, correlation matrices of each cell pair within the ensemble in 5-min. bins. Bottom, circular layout of network graph and clustering coefficient (CC) at each time bin. G) Ensemble clustering coefficients of the fading ensembles across time during Reversal. In all mice that had at least 3 remodeling ensembles (n=5 mice), clustering coefficients of their fading ensembles decreased over time (one-way ANOVAs, all p<0.01). This effect was replicated also by averaging within mice (F_5,20_=44.5, p=7.0e-5).

### Ensembles remodel due to decoupling of weakly-connected neurons

Having observed that ensemble remodeling occurs more frequently during the memory-updating phase of the task, and that their prevalence was correlated with enhanced behavioral performance, we next sought to characterize the relationships between neurons belonging to those ensembles over the course of learning and memory-updating. Based on theories suggesting that ensemble membership may change during learning to prevent overreliance on a single neuronal population^1,47,48^, we hypothesized that the decrease in ensemble activation strength may be a result of certain neurons decoupling from the rest of the ensemble. To investigate this possibility, we used graph theory^49^ to infer and track the remodeled ensembles’ neuron-pair relationships during Reversal. First, we split the Reversal session into 5-min bins. Then in each 5-min bin, for each remodeling ensemble, we correlated the deconvolved calcium activity of all pairs of ensemble member neurons. From these correlation matrices, we constructed undirected graphs where each neuron was a node and edges (connections) were placed between neuron pairs with statistically significant correlations. This yielded one graph per 5-min time bin per remodeled ensemble that collectively described the ensemble’s internal connectivity over time (Fig. 3f). To test whether neurons were decoupling from the rest of the ensemble over Reversal, we computed the average clustering coefficient (a measure of node neighborhood co-activity) in each time bin. In all mice, the average clustering coefficient decreased over time during the Reversal session in remodeling ensembles (Fig. 3g), but not non-remodeling ensembles (Supplementary Fig. 6c). This evidence indicates that hippocampal ensembles remodel due to decoupling of internal co-activity during memory-updating.

If the internal co-activity of remodeling ensembles decreases during memory-updating, which neurons were most likely to be the ones that “dropped”? Previous studies have shown that memories are preferentially stored in neurons that are well-connected with their peers^50–54^. Thus, to update the memory during Reversal, we reasoned that the neurons that dropped from the ensemble may be ones that were relatively weakly co-active with the rest of the ensemble to begin with, but the ones that remained were consistently strongly co-active. To test this possibility, we tracked the degree (i.e., number of connections) of neurons over the course of the Reversal session. First to see whether the connectivity of individual neurons decreased during memory-updating, we computed the average degree of all neurons in remodeled ensembles and confirmed that they decreased over Reversal (Fig. 4a). However, the neuronal population did not homogenously decrease co-activity; certain neurons retained their co-activity while others significantly decreased (Supplementary Fig. 7a-b). Thus, we used the last 5 mins of the Reversal session to identify neurons with high or low co-activity (degree) so that we could then examine whether a neuron’s co-activity at the beginning of the Reversal session might predispose it to be dropped by the end of the session. To do this, we computed the degree of each neuron at the last 5-min time bin of Reversal and partitioned the degree distribution of each remodeled ensemble into quartiles. Dropped neurons were ones whose degree fell in the bottom quartile of the distribution and hub neurons were the ones that fell in the top quartile, strictly at the end of the Reversal session (Fig. 4b-c). We confirmed that dropped neurons significantly decreased their degrees over Reversal, but hub neurons did not (Supplementary Fig. 7a-c). From here, we sought to examine whether a neuron’s degree at the beginning of the Reversal session might predispose it to be either a hub or a dropped neuron at the end of the session. To do this, we compared the degrees of hub and dropped neurons at the *beginning* of the Reversal session and found that within a remodeled ensemble, the dropped neurons had lower degrees than their peer hub neurons, even before they dropped (Fig. 4d). This suggests that the neurons that tend to be dropped have initially lower co-activity with other neurons in the ensemble at the start of the session.

**Figure 4.**
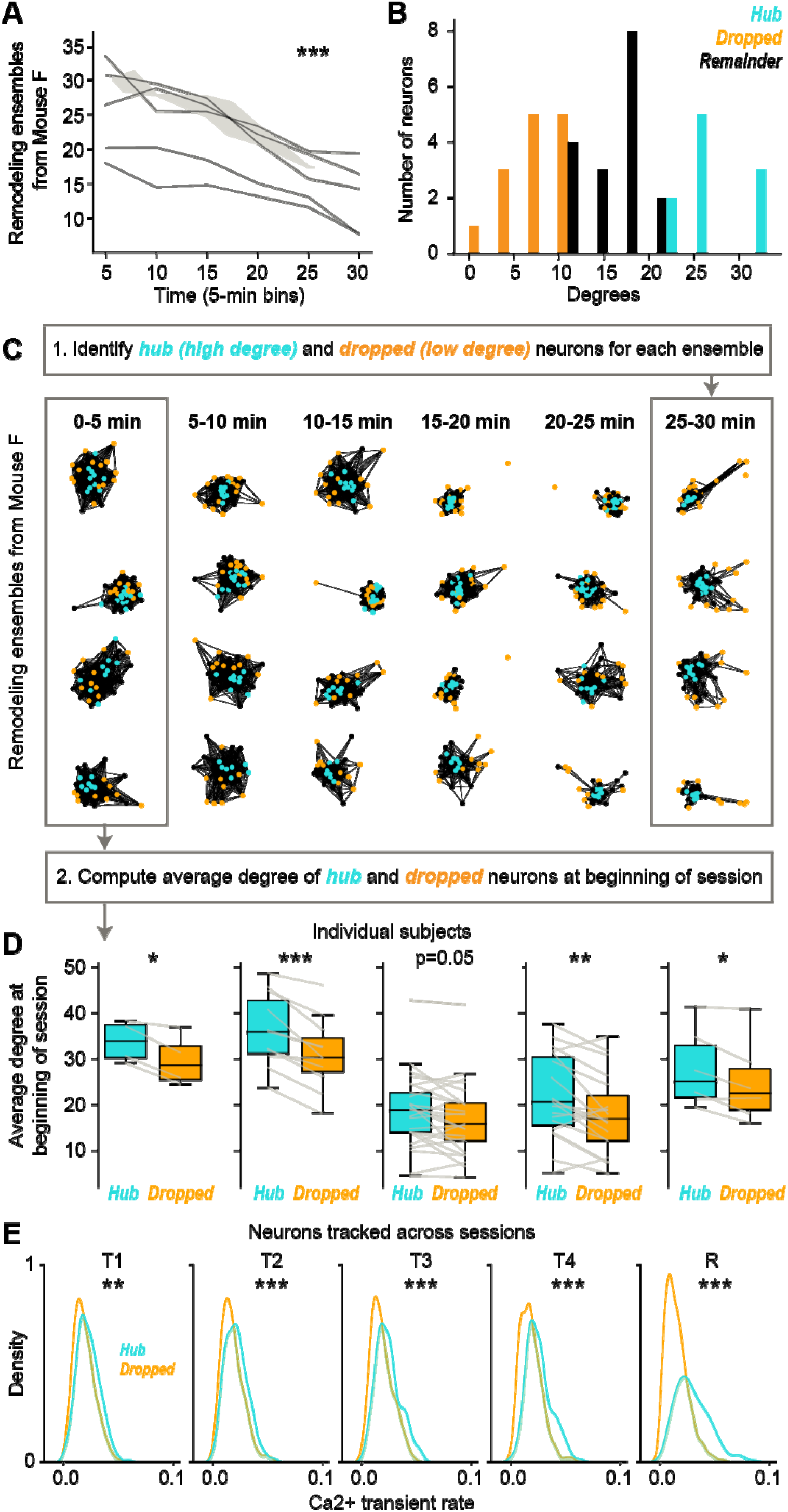
Dropped neurons in remodeled ensembles have low co-activity with other ensemble members at the beginning of the Reversal session. A) Average degrees of the remodeled ensembles across time during Reversal. In all mice that had at least 3 remodeled ensembles (n=5 mice), average degrees of the neurons in their remodeled ensembles decreased over time (one-way ANOVAs, all mice p<0.002). This effect was replicated also by averaging within mice (F_5,20_=27.4, p<0.001). B) Example distribution of degrees from the last 5 min of a remodeling ensemble from Mouse F. We constructed graphs (see Methods) in 5-min time bins on the Reversal session. To identify “dropped” neurons (orange), we found the neurons whose degree fell within the bottom quartile of all degrees from the last 5-min time bin. “Hub” neurons (cyan) were neurons who degree fell within the top quartile. C) Spring layout graphs of remodeled ensembles over 5-min time bins on the Reversal session. Note that the hub neurons reside near the center of the spring projection but the dropped neurons get “pushed” towards the periphery with each time bin. Cyan dots denote hub neurons, orange dots denote dropped neurons and black dots denote dropped neurons.. D) Average degrees of hub versus dropped neurons for each remodeled ensemble and each mouse at the beginning of Reversal. Each line shows the average degree of hub versus dropped neurons within a remodeling ensemble. In all mice, dropped neurons had lower average degrees than hub neurons in their respective ensembles, even before they dropped from the ensemble (all p≤0.05 after Benjamini-Hochberg corrections for multiple comparisons). Analysis repeated with linear mixed models also yielded si^≤^gnificant differences between hub and dropped neurons (F_1,1119.7_=72.6, p<0.0005). E) Event rates of hub and dropped neurons from the end of Reversal, tracked across sessions. Hub neurons in previous sessions had significantly higher calcium event rates compared to dropped neurons, but the difference widened with temporal distance from the Reversal session. Linear mixed models found a main effect of hub versus dropped neurons (neuron type) F_1,1550.4_=130.8, p<0.0005; a session main effect F_4,2936.7_=33.02, p<0.0005; and a neuron type x session interaction F_4,2938.5_=73.48, p<0.0005. Asterisks indicate significance based on paired t-tests on estimated marginal means of the linear mixed model (*p<0.05, **p<0.01, ***p<0.001).

Because the persistence of a neuron within an ensemble can be predicted by early indications of co-activity with its peers, we were curious how far back in time hub and dropped neurons were distinguishable. Previous studies have shown that neuronal excitability was predictive of whether memories were allocated to neurons, but that this mechanism operates within a limited time window^28,55,56^. Therefore, we hypothesized that a similar time-dependent mechanism could influence whether a neuron dropped from an ensemble during future memory-updating. We reasoned that on recent sessions relative to the Reversal session, activity rates may be more predictive of whether a neuron leaves its ensemble compared to sessions farther in the past. We tested this possibility by comparing the calcium transient rates of the Reversal session’s hub and dropped neurons on previous Training sessions. If there was a time-dependent effect of whether activity rates predicted dropping from the ensemble, the difference between the activity rates of hub and dropped neurons should be larger at recent time points and smaller at further time points relative to Reversal. As we expected, neurons classified as hub neurons had higher calcium transient rates compared to their dropped peers, even on the Training sessions days prior, but the difference between hub and dropped neurons’ activity rates decreased the farther back in time we went relative to the Reversal session (Fig. 4e; Supplementary Fig. 8). Overall, these results show that remodeling ensembles expressed during Reversal are made up of neurons that are dissociated from its peers during memory-updating. The neurons that are pruned are ones that are least correlated with their peers, and the least active, even on sessions prior to the pruning.

### Middle-aged mice are impaired at memory-updating and lack the neural mechanisms associated with memory-updating

The data presented thus far suggest that memory-updating is supported by ensemble remodeling where individual neurons within an ensemble become decorrelated within its local network. We predicted that a mouse model that was impaired in spatial memory-updating would also have impaired ensemble remodeling. Therefore, we used aging as a manipulation to see whether we could incur memory-updating deficits alongside ensemble remodeling deficits. Notably, spatial reversal learning is impaired in elderly humans and rodents^6,57,58^. Importantly, previous studies have also shown that middle-age mice are able to learn single hippocampus-dependent memories similar to young adult mice but have difficulties with updating and integrating new information^8,28,59^. To test our hypothesis that impaired ensemble remodeling leads to deficits in memory-updating, we used middle-aged mice (16–19-month old), who were able to learn the initial goal locations just as well as young adult (6-10-month old) mice (Fig. 5a). However, they were significantly impaired during Reversal, showing a worse correct rejection rate than young mice (Fig. 5b). Middle-aged mice did not differ in the number of trials ran nor number of rewards received compared to young mice (Supplementary Fig. 9a-c). As predicted, they failed to show an increase in the proportion of remodeling ensembles during Reversal, compared to the young mice (Fig. 5c). Similar to young adult mice, the proportion of remodeling ensembles in middle-aged mice correlated with behavioral performance (Fig. 5d), suggesting that this ensemble remodeling is related to memory-updating. Together, these data indicate that middle-aged mice are impaired at memory-updating and they lack the ensemble remodeling mechanisms that are normally seen in earlier adulthood.

**Figure 5.**
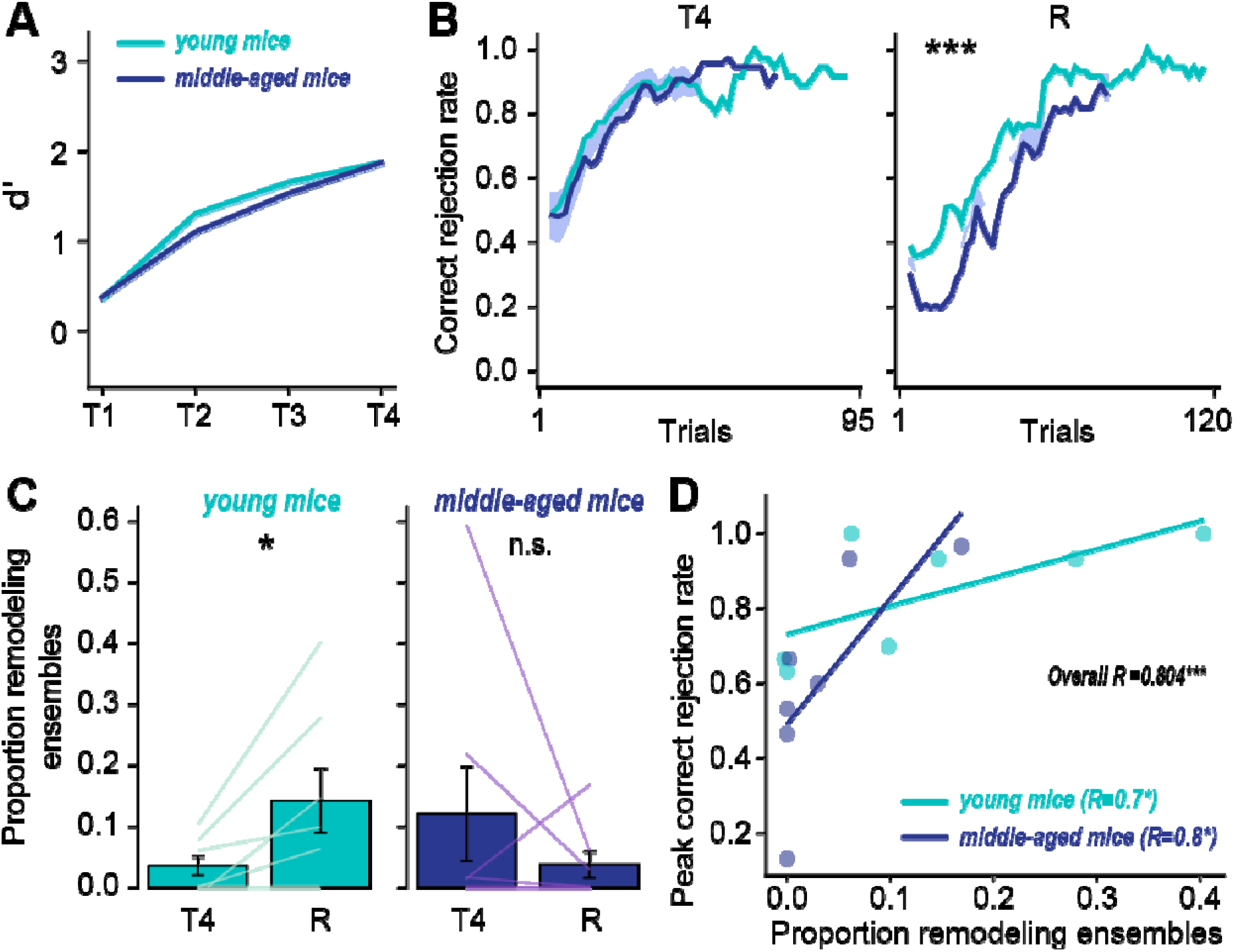
Middle-aged mice exhibited impairments in memory-updating and failed to show normal ensemble flexibility that was seen in young adult mice. A) Behavioral d’ in young and middle-aged mice. Both groups learned the task (main effect of session with repeated-measures ANOVA F_3,39_=39.3, p<0.0001), but there was no significant difference between age groups (F_1,13_=-0.148, p=1.0). Training1-4 abbreviated as T1-4. Young mice data was duplicated from Fig. 1B. B) Correct rejection rate of young compared to middle-aged mice over trials on Training4 (T4) and Reversal (R). During Reversal, there was a significant effect of trials (two-way ANOVA F_1,57_=14.8, p<0.0001) and between young and middle-aged mice (F_1,57_=90.0, p<0.0001). On Training4, there were no differences between the groups (F_1,64_=2.17, p=0.14). C) Proportion of remodeling ensembles during Training4 compared to Reversal. There were no significant differences between the two sessions in middle-aged mice (Wilcoxon signed-ranked test W=8.0, p=0.31). Young mice data duplicated from Fig. 3C. D) Scatter plot of proportion of remodeling ensembles against peak correct rejection rate during Reversal. There was a significant correlation between the proportion of remodeling ensembles and correct rejection rate in middle-aged mice (Spearman R=0.808, p=0.014). Young mice data duplicated from Fig. 3D. There was a significant correlation even pooling the two age groups (Spearman R=0.804, p<0.001)

## Discussion

### Ensemble remodeling enables adaptive memory-updating and forgetting

Forgetting is functionally important because deconstructing previous memories allows the organism to behave more flexibly^4^. Here, we subjected mice to a paradigm where forgetting previous reward locations would be beneficial for efficient reward retrieval. They responded with memory-updating and ensemble remodeling (Fig. 6). To our knowledge, our study was the first to show that hippocampal ensembles undergo remodeling as mice underwent memory-updating. Reversal learning has been predominantly attributed to the frontal cortices^20,60–63^, so our study on the hippocampal contribution to this process adds new insight on how this region could support memory flexibility. Given the known relationship between the hippocampus and frontal cortices during memory consolidation^64^, it is highly likely that successful memory-updating depends on direct interplay between these two regions. Future work could examine how downstream regions interact with remodeled hippocampal activity.

**Figure 6.**
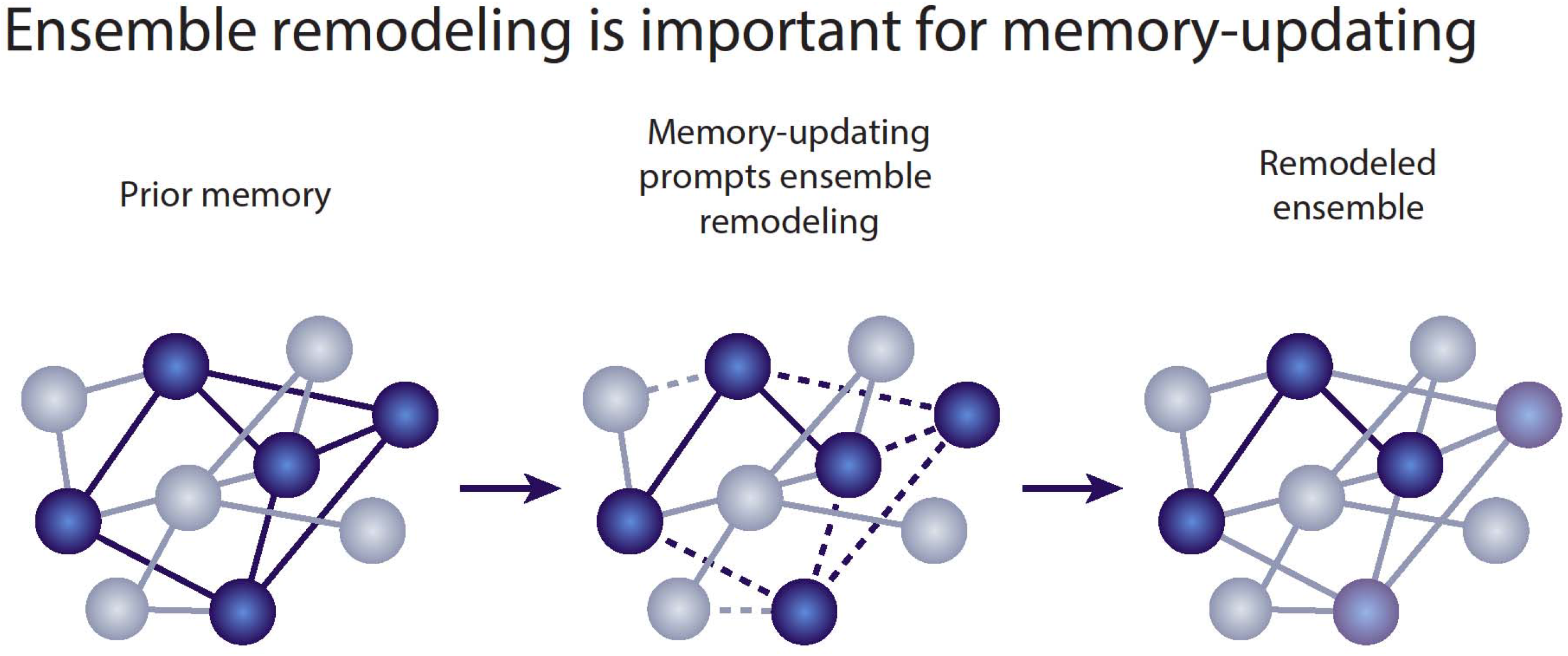
Schematic of ensemble remodeling during memory-updating. A well-learned memory is encoded in a neuronal ensemble (blue spheres in the left diagram). During memory-updating, when environmental stimuli fail to match expectations based on prior experience, memory-updating prompts the ensemble to undergo remodeling (center). The remodeled ensemble reflects the updated memory (right).

The breakdown of hippocampal ensembles in our task appears to be vital for the network to form new functional connections and update an already existing memory. Ensemble remodeling may address the problem of memory capacity in the brain that comes with continual learning. If memories persisted in the same ensembles indefinitely, the brain would not contain enough neurons to store more and more new information as an animal learns over its lifetime^3^. Instead, ensembles must be remodeled to assimilate new information. Aside from storage requirements, ensemble remodeling may be adaptive by promoting network flexibility. Dropout of artificial neurons is sometimes employed in artificial neural networks to prevent overfitting^48^, which reduces reliance of the network (*in silico* or *in vivo*) on an overburdened set of neurons. This has the effect of a more robust, distributed network for learning new information. Accordingly, artificial neural networks have been found to better learn conflictual information (e.g., reversal learning) when flexibility is built into those networks^65,66^.

### Aging as a model of impaired memory-updating and ensemble remodeling

Rather than using neurotechnology to artificially manipulate ensemble dynamics, we chose to use middle-aged mice to investigate how natural deterioration of ensemble remodeling might occur alongside natural deterioration of memory flexibility. Although this approach comes with the burden of aging covariates, there is an overwhelming benefit to being able to directly tie our findings to translational models. Moreover, our findings reflect naturalistic neurobiological activity that cannot be attributed to off-target effects of artificial manipulation. In our middle-aged cohort, we found that although remodeled ensembles were rare (Fig. 5c), their prevalence still correlated with behavior (Fig. 5d). This suggests that certain privileged middle-aged mice that can harness the neuroplasticity required to remodel ensembles may have unique advantages in memory-updating. That idea is consistent with past findings showing that although most aged rodents fail to learn trace eyeblink conditioning, some do succeed and those that succeeded showed normal hippocampal neurophysiological properties (e.g., neuronal excitability), similar to their younger counterparts^67–70^. The common ground between these studies and ours is that aging broadly affects memory performance, but resilient middle-aged individuals with intact hippocampal mechanisms for learning can perform at young-like levels. Such a finding has important clinical implications that may allow us to better understand how to treat poor memory and cognitive flexibility in aged humans^5,6^.

### Possible mechanisms underlying ensemble remodeling

Mechanistically, ensemble remodeling might be achieved through synaptic plasticity. More specifically, the rearrangement of synaptic weights might disfavor certain neurons such that they no longer become co-activated with the rest of the ensemble (i.e., get dropped). This is consistent with previous studies reporting reductions in dendritic spine size and density in engram neurons as their encoded memories faded^71–73^. Also in support of this neuroplasticity hypothesis, blocking hippocampal LTD impairs spatial reversal learning^25,26^, possibly by disrupting ensemble remodeling. However, future technologies that can measure synaptic strengths at the population level would be needed to test whether the remodeling ensembles we observed are indeed undergoing LTD. We further speculate that breakdown of ensembles during memory-updating may be NMDA receptor-dependent in the hippocampus, as two previous studies have described NMDA receptor-mediated LTD underlying memory flexibility^25,27^. Another key piece of evidence is that hippocampal spatial representations are paradoxically more stable when NMDA receptors are antagonized^74^, in spite of their canonical role in memory stabilization^75,76^. Those findings suggest that NMDA receptors may play an understudied role in ensemble disassembly during memory-updating.

Ensemble remodeling could also result from ramping up inhibition onto the dropped ensemble member neurons, perhaps in concert with the slower LTD mechanisms. Interneurons have been shown to constrain ensemble size in the hippocampus^77^, done by exerting inhibitory influence onto a select group of principal cells in order to maintain homeostatic balance across the network. Such a mechanism may be important to prevent runaway excitation when ensembles are forced to remodel during memory-updating events. This inhibition could be mediated by highly specific interneuron engram networks^78–80^. Future work could test whether local hippocampal interneurons are enforcing a competitive, lateral inhibition-like regime such as in the amygdala^55^ to suppress disfavored neurons during memory-updating. These suppressed excitatory neurons would then drop out of the ensemble to allow room for a modified memory representation. The activity changes brought about by this inhibition may then result in long-lasting changes in synaptic strength that persist over longer time periods beyond that of the immediate memory-updating episode.

### Heterogeneous remodeling of hippocampal ensembles

In our graph theory analyses, we found that neurons that will prospectively drop from ensembles during memory-updating were the least connected with the rest of their ensemble member peers to begin with. This finding is consistent with others showing that ensemble maintenance is heterogenous^81^ and not random, favoring the neurons that have high functional importance and pruning those that have lesser significance. For example, Arc-expressing neurons show more synaptic stability and are more integrated in memory ensemble networks^53^. Neurons that have high selectivity for specific spatial trajectories are also selectively stabilized during navigational memory tasks^82,83^. During hippocampal sharp-wave ripples, synapses are potentiated^84^, but synapses in non-engram neurons undergo LTD^85^, suggesting that the less critical neurons are purged from the ensemble. Memory-updating during our task may be engaging inhibition, neuroplasticity, or both mechanisms to reduce the engagement of dropped neurons, perhaps with the endpoint of enhancing signal-to-noise. Though our study makes considerable advances in understanding the neural mechanisms of memory-updating, further study is needed to fully grasp the whole-brain dynamics that underlie flexible memory.

## Acknowledgements

We thank Daniel Aharoni and his lab for their tremendous work in developing the UCLA V4 Miniscope. We thank Caitlin Vander Weele, Gabi Serrato Marks, and Stellate Communications for their help in figure editing. We thank Yaniv Ziv, Priya Rajasethupathy, Lisa Giocomo, Erin Rich, Peter Rudebeck and their lab members for helpful comments on the preliminary results of this manuscript. We thank Paul Philipsberg for his assistance in designing the experimental apparatus. We also thank Alexa LaBanca, Yosif (Joe) Zaki, Brian Sweis, Austin Bagetta, Yu (Susie) Feng, Zoé Christenson-Wick, Lingxuan Chen, and Iván Soler for helpful comments throughout this project. W.M. was funded by F32AG067640. T.S. was funded by the CURE Epilepsy Taking Flight Award, American Epilepsy Society Junior Investigator Award, R03 NS111493, R21 DA049568, and R01 NS116357. D.J.C was funded by NIH DP2MH122399, R01 MH120162, One Mind Otsuka Rising Star Award, McKnight Memory and Cognitive Disorders Award, Klingenstein-Simons Fellowship Award in Neuroscience, Mount Sinai Distinguished Scholar Award, Friedman Brain Institute Scholar Award, Brain Research Foundation Award, Irma T. Hirschl/Monique Weill-Caulier Research Award and NARSAD Young Investigator Award.

## Methods

### Animals

All protocols were approved by the Institutional Animal Care and Use committee at the Icahn School of Medicine at Mount Sinai, in accordance with National Institutes of Health guidelines. A total of 31 adult C57BL/6 mice of both sexes, from the National Institute on Aging, were used in our experiments. We used 17 of these mice (5-6 months old) in our PSAM experiment and 14 mice in our calcium imaging experiment. Mice were group-housed with littermates on a 12 h light/dark cycle until behavioral experiments began, whereupon they were single-housed. Mice were randomly assigned to receive either PSEM or vehicle injections during Reversal. Young adult mice were 6-10 months old and aged mice were 16-19 months old.

### Viral constructs

For our PSAM virus, we used AAV5-syn-PSAM^4^-GlyR-IRES-eGFP (Addgene viral prep # m119742-AAV5; http://n2t.net/addgene:119742; RRID:Addgene_119742) with a titer of ∼2.4 x 10^13^ GC/mL. For our calcium indicator virus, we used AAV1-syn-GCaMP6f-WPRE-SV40 (Addgene viral prep # 100837-AAV1; http://n2t.net/addgene:100837; RRID:Addgene_100837) diluted to a titer of ∼5.0 x 10^12^ GC/mL with phosphate buffered saline.

### Drugs

We used uPSEM-817 (Tocris) as our ligand to inactivate PSAM-expressing hippocampal neurons, diluted in saline to a concentration of 0.1 mg mL^-1^, corresponding to a 1 mg kg^-1^ dose per mouse. PSEM or vehicle (saline) was injected intraperitoneally 30 min before behavior, corresponding to the time required for PSEM to cross the blood brain barrier^33^.

### Apparatus

We used two environments for mice to navigate during testing. Wallpapers were placed on the outside and inside walls of the environments to provide local spatial cues, while lamps and furniture were arranged in the room to provide distal cues. Mice were able to see distal cues as they would frequently rear and place their front paws on the tops of the track walls to scan their surroundings.

One environment was a 100 cm long x 6.3 cm wide x 10 cm high linear track fabricated from white acrylic (Maze Engineers). At each end of the linear track were blunted 18 g needles, connected to a gravity-assisted water dispensing system controlled by an Arduino Uno. Water was delivered when a touch sensor circuit detected a lick on either port.

The second environment was a 6.3 cm wide x 10 cm high circular track with a diameter of 76.2 cm, also made of white acrylic (Maze Engineers). Eight water ports were arranged on the outside wall of the track every 45 degrees along the circular track. For calcium imaging experiments, we used a commutator (Neurotek, DragonFly) to prevent cable tangling during the behavior. Miniscope recording, behavioral recording, and lick detection were all triggered simultaneously using a 3.3V digital signal from the Arduino Uno using a custom Python script.

### Circular track spatial reversal task

For all experiments, mice were handled for 5 min for at least two days prior to behavioral testing. Mice were then water-deprived and body weight was monitored daily to ensure that values did not drop below 85% of their starting weight. Prior to testing on the circular track, mice were habituated to licking for water rewards on the linear track for 5-7 days for 20 min each day. Mice had to retrieve water at both port locations. An air filtration fan was turned on for the entire duration of all testing days for each experiment to provide white noise. Audible clicks played when water was delivered.

Following linear track habituation, mice were trained to retrieve water rewards at two pseudorandom reward ports on the circular track (Training1-4). The reward ports had to be separated by at least two other ports to prevent the mouse from simply shuttling between two adjacent ports without traversing the entire track. The mice were forced to navigate only in the clockwise direction.

Sessions were 30 min long. On the Reversal day, we selected two new pseudorandom reward ports with the same spacing requirement and an additional requirement that the new reward ports could not overlap with the previous two reward ports. There were no indications given to the mouse that the reward locations had changed.

### Stereotaxic surgeries

Mice were anesthetized with 1-2% vaporized isoflurane and placed into a stereotaxic frame (Kopf). For PSAM experiments, mice received a bilateral infusion of 400 nL AAV5-syn-PSAM^4^-GlyR-IRES-eGFP into dorsal CA1 (AP -2.0 mm relative to Bregma, ML ±1.5 mm, DV -1.5 mm) using a Nanoject microinjector (Drummond Scientific) at a rate of 2 nL s^-1^. Sutures were applied and mice were allowed to recover in a heated cage. Mice received subcutaneous injections of dexamethasone, ampicillin, and carprofen daily for 7 days post-surgery.

For calcium imaging experiments, mice instead received at a unilateral (right hemisphere) infusion of 400 nL AAV1-syn-GCaMP6f-WPRE-SV40 into dorsal CA1 (same coordinates as above). Ten min after viral infusion, they were also given a GRIN lens implant. A ∼2 mm diameter circular craniotomy was opened centered around AP -2.25 mm ML +1.5 mm and the overlying cortex was aspirated until vertical white fiber tracts were visible^86^. Bleeding was controlled with constant irrigation using cortex buffer solution and Surgifoam (Ethicon). Once bleeding had ceased, the GRIN lens (1 mm diameter, 4 mm length, Inscopix) was slowly inserted to the depth of DV -1.3 mm. The gap between the lens and the skull was filled using Kwik-Sil (World Precision Instruments), superglued, and then reinforced with dental acrylic (Ortho-Jet). The top of the GRIN lens was then covered with a layer of Kwik-Cast (World Precision Instruments) and a final layer of dental acrylic for protection until base plate attachment. Four weeks later, we attached inhouse-machined aluminum base plates to permit attachment of the V4 Miniscope, a modified version of the UCLA Miniscope described previously^28,29^. More information on the V4 Miniscope can be found on the project GitHub page: https://github.com/Aharoni-Lab/Miniscope-v4. While mice were anesthetized, the protective acrylic cap over the GRIN lens was drilled off. The base plate was then screwed onto the Miniscope and lowered onto the desired focal plane that best captured the neuronal field of view. The base plate was then superglued and reinforced with dental acrylic. An inhouse-machined Delrin cap was then screwed over the base plate to prevent debris accumulation inside.

### Histology

Mice were anesthetized with isoflurane and transcardially perfused with 4% paraformaldehyde (PFA). Brains were extracted and stored in PFA. Brains sliced using the cryostat were transferred to a 30% sucrose solution for ∼3 days. Slices of 50 µm thickness were acquired using either cryostat or vibratome and mounted with Vectashield Hardset mounting medium with DAPI (Vector Laboratories). Viral injection sites were verified postmortem using a wide field epifluorescent microscope (Leica) with the green channel.

### Behavior tracking and calcium imaging analysis

To track the location of mice for all experiments, we used a combination of ezTrack^87^ and DeepLabCut^88,89^. Positional data was manually curated using custom Python scripts on frames where position was misidentified.

Calcium imaging data was passed through the Minian pipeline to extract neuronal footprints and traces^90^. Minian is a Python package for Miniscope imaging source extraction that utilizes constrained nonnegative matrix factorization (CNMF)^91,92^ to denoise and demix one-photon calcium imaging data. For each mouse, we used the visualization tools of Minian to curate the algorithm parameters that would produce optimal calcium traces and neuronal footprints. Parameters were kept constant across sessions for each mouse. Calcium traces and deconvolved spikes were then synchronized with the behavioral video and lick timestamps using custom Python scripts.

After extracting neuronal footprints and traces, we registered neurons across sessions using the Matlab package CellReg^93^. Minian outputs were converted into Matlab-readable matrices using a custom Python script and passed through CellReg before being converted back into a Python-readable data structure.

### Behavioral analyses

Behavioral performance on the circle track was computed using hit rates, correct rejection rates, and d’. The hit rate per trial was defined as the number of times a rewarded port was licked at least once on that trial divided by the number of rewarded ports (2 ports). Similarly, the correct rejection rate was defined as the number of times an unrewarded port was not licked at all on that trial divided by the number of unrewarded ports (6 ports). To calculate d’, we used the loglinear approach^94^ to prevent infinite d’ rates under edge cases and then followed the formula:

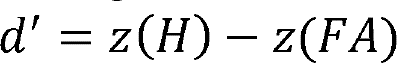

Where H is the loglinear transformed hit rate, FA is the false alarm rate (1 minus the correct rejection rate) and z(x) is the z-transform based on a standard normal distribution.

### Place cell criteria

Spatial rate maps were computed using 2-cm-wide spatial bins and a speed threshold of greater than 7 cm s^-1^ Spatial rate maps were then normalized by the occupancy of the mouse for each spatial bin. Spatial information for each neuron was calculated using the following formula:

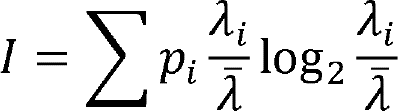

Where p_i_ is the probability of occupancy in the i-th spatial bin (occupancy divided by total recording time), λ_i_ is the calcium activity rate in the i-th spatial bin, 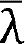 and is the mean calcium activity rate of that neuron. The spatial information for each neuron was then z-transformed relative to a surrogate distribution, a procedure that has been found to more accurately convey spatial information relative to chance^95^. To create the surrogate distribution, we circularly shuffled the spatial position of the mouse relative to the calcium activity and recomputed the spatial information, repeated 500 times. A neuron was classified as a place cell if (1) it had a statistically higher spatial information value as expected from chance, compared to the surrogate distribution (p < 0.001), and (2) it fired at least 40 calcium transients.

### Rate remapping

Rate remapping scores were calculated similar to previous methods in order to determine the extent to which activity rates differed across time in a neuron’s place field^38^. We split each recording session in half and calculated each neuron’s mean activity rates of place fields independently for each half. We determined the place fields of that neuron to be spatial bins where the average activity exceeded 90% peak activity across all spatial bins. Then, the rate remapping score is calculated as the absolute difference of the mean activities across place fields between first and second half of the session, divided by the sum of the mean activities. This yields a range of possible scores from 0 (no rate change) to an asymptotic value of 1 (infinite rate change).

### Ensemble detection and remodeling ensemble classification

Ensembles for each session were detected using a Python implementation of an unsupervised statistical framework based on principal component analysis (PCA) followed by independent component analysis (ICA)^43^. Binarized calcium activity was smoothed using a 5-bin moving average (∼300 ms, slightly larger than a single theta cycle) and z-scored. Using this activity matrix, we performed PCA and compared the eigenvalues of each principal component to a surrogate distribution of eigenvalues, created using activity matrices where all neurons had their activities circularly shuffled relative to each other, repeated 500 times. The number of ensembles (independent components) was the number of principal components that had eigenvalues higher than 99% (p < 0.01) of the eigenvalues from the surrogate distribution. We then use this number as the number of independent components to be extracted with fast-ICA algorithm. Each ensemble (independent component) has a weight vector representing the contribution of each neuron to the ensemble, and neurons were considered ensemble members if their weight exceeded 2 standard deviations above the mean of that ensemble^43,96^. We then used the outer product of the weight vector to produce the projection matrix *P*, which can be used to project neural activity into ensemble activation strength. To compute the activation strength of each ensemble at each temporal bin *b*, we used the formula:

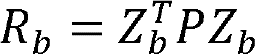

Where *Z_b_*is the normalized calcium activity at time bin *b*, and *P* is the projection matrix, with the main diagonal set to zero to ensure that calcium activity of a single neuron does not contribute to the ensemble strength *R_b_*.

Ensembles were classified as “remodeled” using the non-parametric Mann-Kendall trend test^97^. For an ensemble to be called a remodeled ensemble, its activation strength needed to have a negative slope across time and *p*<0.005 after Šidák correction for multiple comparisons across all ensembles.

### Spatial and lick decoders

Spatial decoders were constructed using naïve Bayes classifiers^98^ trained on 2-cm-wide binned spatial position and calcium activity from all recorded neurons from data where the mouse was moving above the speed threshold of 7 cm s^-1^. Accuracy was assessed using 5-fold cross validation. Chance levels were computed using data where the spatial position of the mouse was circularly shuffled relative to the calcium activity, repeated 100 times.

For decoders that were trained and tested across two sessions with ensemble activity, we registered ensembles using cosine similarities between weight vectors of ensemble pairs. First, we sorted weights for each neuron by their indices based on the registration maps from CellReg. Neurons that were not registered across the two sessions were not included. Then, we computed cosine similarities between each ensemble pair’s weight vectors. Two ensembles were registered if (1) they were mutually most similar to each other (highest cosine similarity) among all other possible pairs and (2) their z-scored cosine similarity exceeded 2.58 (*p*<0.01).

Lick decoders were constructed from random forest classifiers (*n*=100 trees) trained on the ensemble activity of all registered ensembles at each lick. Only the first lick of each trial was considered. Since mice typically licked more at rewarded ports than unrewarded ports, our class labels (reward port identity) were imbalanced. To correct for this, weights for each class were balanced such that the weights were inversely proportional to the frequency at which each reward port was licked, penalizing misclassification of rarely licked ports. The lick decoders were trained on all the data from one session and then tested on all the data from the second session. Chance was computed using accuracy measurements from classifiers trained on data where the labels (reward port identity) were shuffled.

### Ensemble graphs

Ensemble graphs^99^ were constructed from pairwise Spearman correlations between ensemble member calcium activities (Fig. 3-5). A neuron was considered an ensemble member if its weight exceeded 2 standard deviations above the mean of all weights. We split sessions into six 5-min time bins, then computed the Spearman correlation between each ensemble members’ calcium activities for each time bin. Edges (connections) were assigned between nodes (ensemble member neurons) if their correlation p-value was less than 0.05 after false discovery rate corrections for multiple comparisons. The clustering coefficient for each node was defined as:

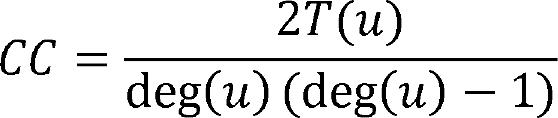

Where T(u) is the number of triangles through node u and deg(u) is the degree (number of connections) of node u.

We classified hub and dropped neurons based on their degrees (Fig. 4). We first split the data into 5-min time bins. Then, for each remodeled ensemble, we computed the degrees of all its member neurons separately for each time bin. In the last 5-min time bin, we divided the distribution of degrees across all neurons into quartiles. Neurons were classified as hub neurons if their degrees were in the highest quartile. Neurons were classified as dropped neurons if their degrees were in the lowest quartile. For each neuron, we then took the difference of degrees between the last and first time bin to calculate the extent to which connections decreased across Reversal (Fig. 4c). We also examined the degrees at the first time bin and compared hub versus dropped neurons to assess whether hub and dropped neurons differed in their degrees at the beginning of Reversal (Fig. 4d).

### Statistics

Statistics were performed using Python statistics libraries including pyMannKendall^97^, sklearn^98^, networkx^99^, scipy^100^, numpy^101^, pandas^102^, and pingouin^103^. Some analyses were also performed with R, using lme4^104^ and lmerTest^105^ to compute linear mixed models on calcium transient rates and degree with neuron category (hub or dropped) and session (if appropriate) as fixed effects and mouse, ensemble nested within mouse, and neuron nested within mouse as random effects. Statistical significance was measured using two-tailed Wilcoxon signed-rank tests, paired and unpaired t-tests, one- or two-way analysis of variance (ANOVAs) with interaction terms, chi square tests, Mann-Kendall trend tests, and Spearman correlations, unless otherwise noted. Post-hoc tests on linear mixed models were done with pairwise t-tests using estimated marginal means (emmeans package in R^106^). P-values (*P<0.05, **P<0.01, ***P<0.001) were corrected for multiple comparisons using Sidak’s or Benjamini-Hochberg false discovery rate adjustments. In boxplots, the line represents the median, the box represents the1^st^ and 3^rd^ quartiles, and the whiskers represent the data range. Sample sizes were not predetermined, but were comparable to previous publications^29,107^. In our PSAM experiments, 1 mouse was dropped due to data corruption, 4 due to viral mistargets, and 2 due to inability to learn the task (predefined as d’<1 on the Training4 session), leaving a total of n=17 mice in that experiment.

---

**Supplementary Table 1.** PSEM vs vehicle 2-way ANOVA results on behavioral performance after downsampling trials to match the lowest number of trials across all mice in Training4 and Reversal. PSEM mice were impaired on all metrics even with trial downsampling.

| Test | F statistic | p-value |
| --- | --- | --- |
| d': PSEM vs vehicle on Training4, after downsampling to match trial counts | Main effect of trial F=0.96 | 0.42 |
|  | Main effect of group F=0.54 | 0.48 |
|  | Trial x group interaction F=0.34 | 0.79 |
| d': PSEM vs vehicle on Reversal, after downsampling to match trial counts | Main effect of trial F=1.60 | 0.20 |
|  | Main effect of group F=11.6 | 0.0012** |
|  | Trial x group interaction F=1.87 | 0.14 |
| Hit rate: PSEM vs vehicle on Training4, after downsampling to match trial counts | Main effect of trial F=0.13 | 0.94 |
|  | Main effect of group F=1.16 | 0.29 |
|  | Trial x group interaction F=0.38 | 0.76 |
| Hit rate: PSEM vs vehicle on Reversal, after downsampling to match trial counts | Main effect of trial F=1.75 | 0.17 |
|  | Main effect of group F=22.1 | 0.00002*** |
|  | Trial x group interaction F=0.40 | 0.75 |
| Correct rejection rate: PSEM vs vehicle on Training4, after downsampling to match trial counts | Main effect of trial F=1.26 | 0.30 |
|  | Main effect of group F=0.22 | 0.64 |
|  | Trial x group interaction F=0.08 | 0.97 |
| Correct rejection rate: PSEM vs vehicle on Reversal, after downsampling to match trial counts | Main effect of trial F=0.71 | 0.55 |
|  | Main effect of group F=6.24 | 0.015* |
|  | Trial x group interaction F=0.30 | 0.83 |

**Supplemental Figure 1.**
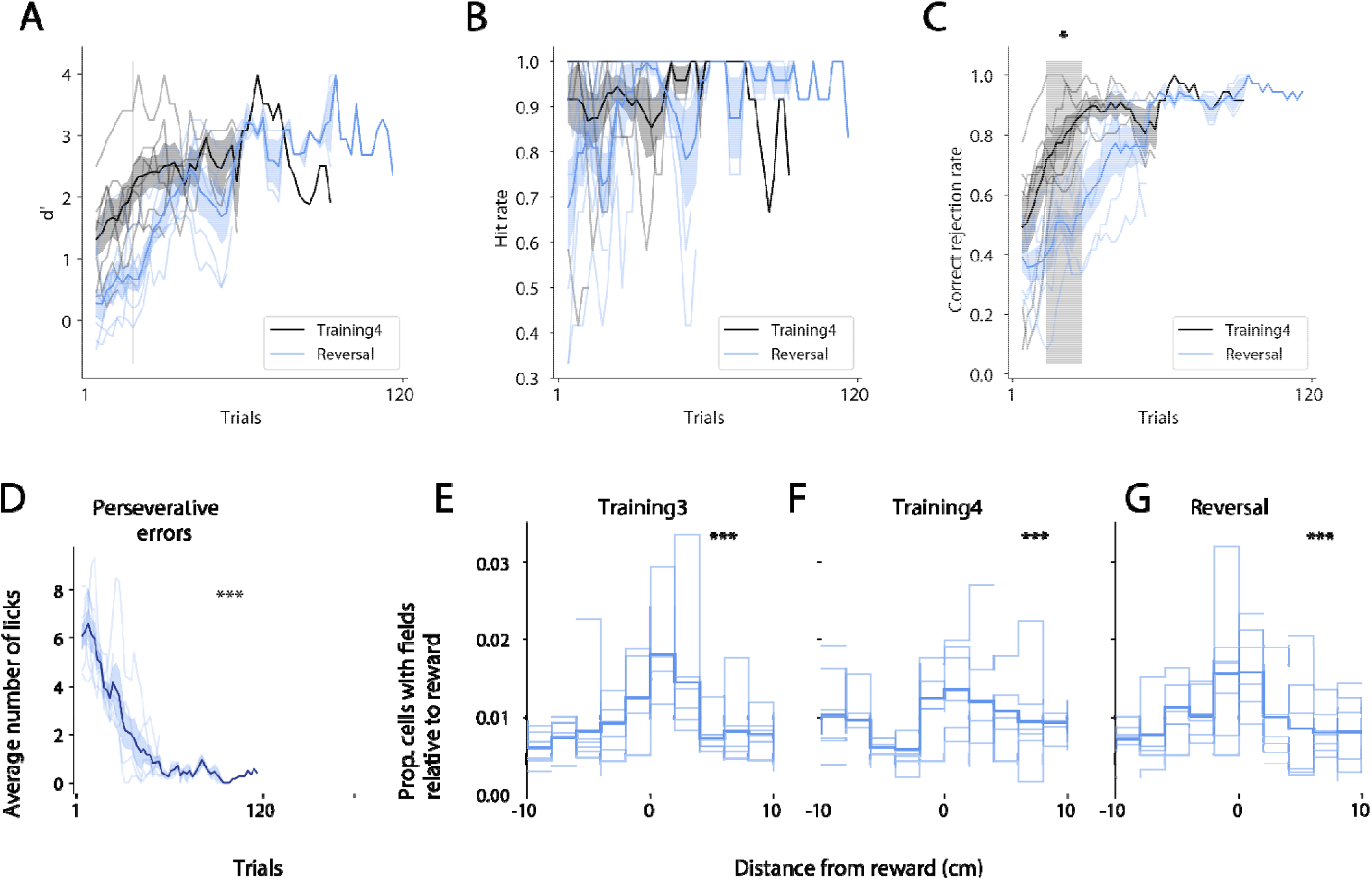
Behavior and place field density of calcium imaging mice. A) Behavioral d’ of calcium imaging mice on Training4 versus Reversal. B) Hit rate on Training4 versus Reversal. C) Correct rejection rate on Training4 versus Reversal. Gray shaded region indicates significant differences across the two sessions (paired t-tests, p<0.05 after Sidak correction for multiple comparisons) D) Perseverative errors over trials during Reversal. Errors significantly decreased over time (linear mixed models with mouse ID as a random effect, F_1,199.77_=205.8, p<0.0001). E) Distribution of spatial field centers relative to reward locations on Training3. Light lines represent individual mice. Thick line represents probability distribution across all mice. There was an overrepresentation of reward sites (chi square test against a uniform distribution, p<0.0001). F) Distribution of spatial field centers for Training4 (chi square test, p<0.0001). G) Distribution of spatial field centers for Reversal (chi square test, p<0.0001).

**Supplemental Figure 2.**
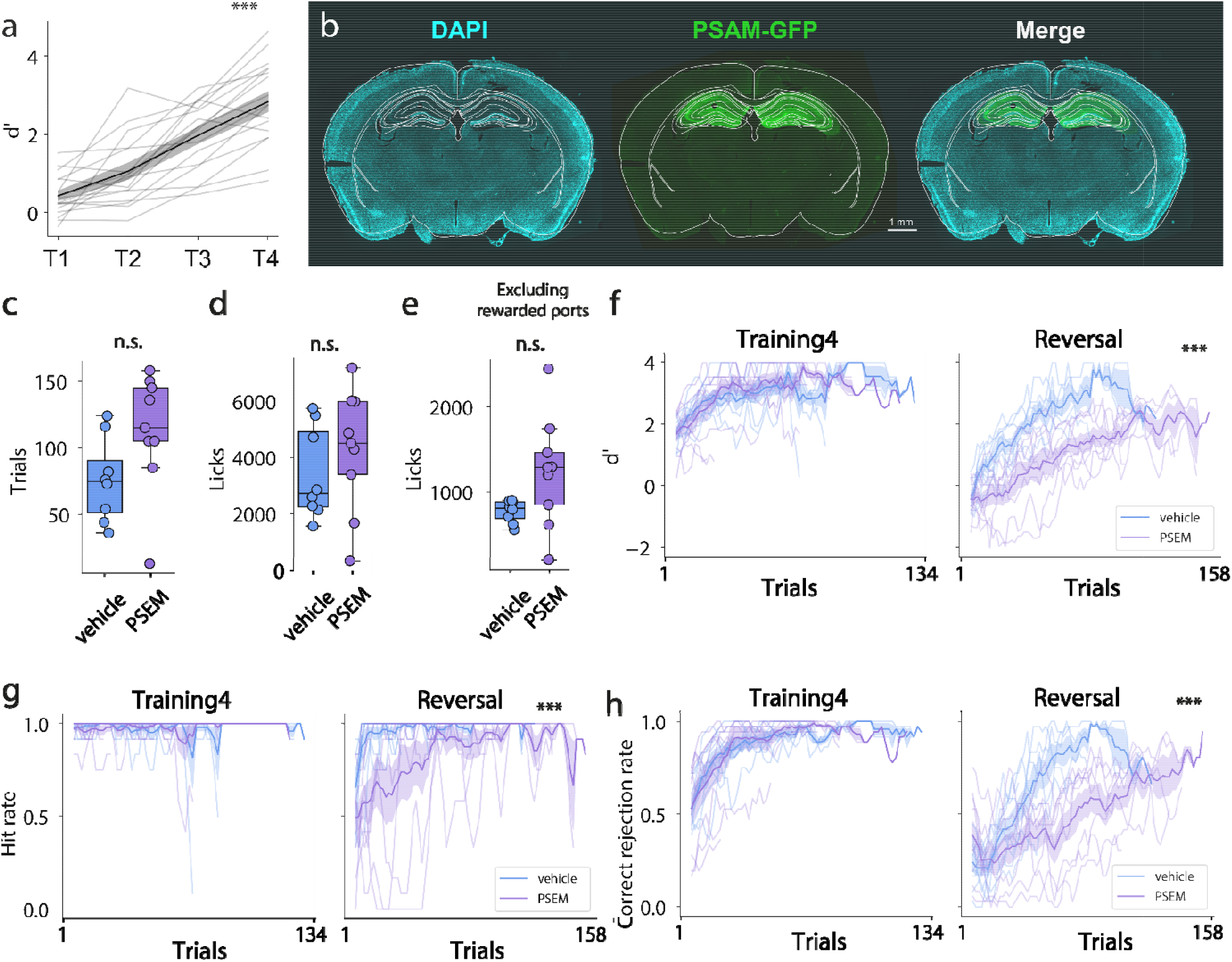
The dorsal hippocampus is required for spatial reversal. A) Behavioral d’ of PSAM cohort over Training. Mice improved their performance over Training (repeated-measures ANOVA F_3,48_=60.7, p<0.0001). B) Example coronal slice stained for PSAM-GFP and DAPI. C) Trial counts in vehicle and PSEM mice during Reversal. There was no significant difference between groups (t_15_=-1.94, p=0.07). D) Number of licks during Reversal. There was no significant difference between groups (t_15_=-0.87, p=0.40). E) Number of licks during Reversal, excluding rewarded ports. There was no significant difference between groups (t_15_=-1.97, p=0.07). F) Behavioral d’ of vehicle and PSEM groups during Training4 and Reversal (PSEM mice did not receive PSEM during Training4). There was a significant difference between PSEM and vehicle during Reversal, but not Training4 (Training4: two-way ANOVA main effect of trial F_1,64_=4.41, p<0.001; no main effect of group F_1,64_=0.90, p=0.34; no group x trial interaction F_1,64_=0.72, p=0.95; Reversal: main effect of trial F_1,76_=8.58, p<0.001; main effect of group F_1,76_=418.7, p<0.001; no group x trial interaction F_1,76_=1.05, p=0.37). G) Hit rate during Training4 and Reversal. There was a significant difference between PSEM and vehicle during Reversal, but not Training4 (Training4: two-way ANOVA no main effect of trial F_1,64_=0.72, p=0.95; no main effect of group F_1,64_=0.0, p=0.97; no group x trial interaction F_1,64_=0.52, p=0.99; Reversal: main effect of trial F_1,76_=1.85, p<0.001; main effect of group F_1,76_=114.4, p<0.001 no group x trial interaction F_1,76_=0.70, p=0.97). H) Correct rejection rate during Training4 and Reversal. There was a significant difference between PSEM and vehicle during Reversal, but not Training4 (Training4: two-way ANOVA main effect of trial F_1,64_=6.64, p<0.001; no main effect of group F_1,64_=0.06, p=0.81; no group x trial interaction F_1,64_=0.32, p=0.99; Reversal: main effect of trial F_1,76_=6.54, p<0.001; main effect of group F_1,76_=175.0, p<0.001; and group x trial interaction F_1,76_=1.94, p<0.001).

**Supplementary Figure 3.**
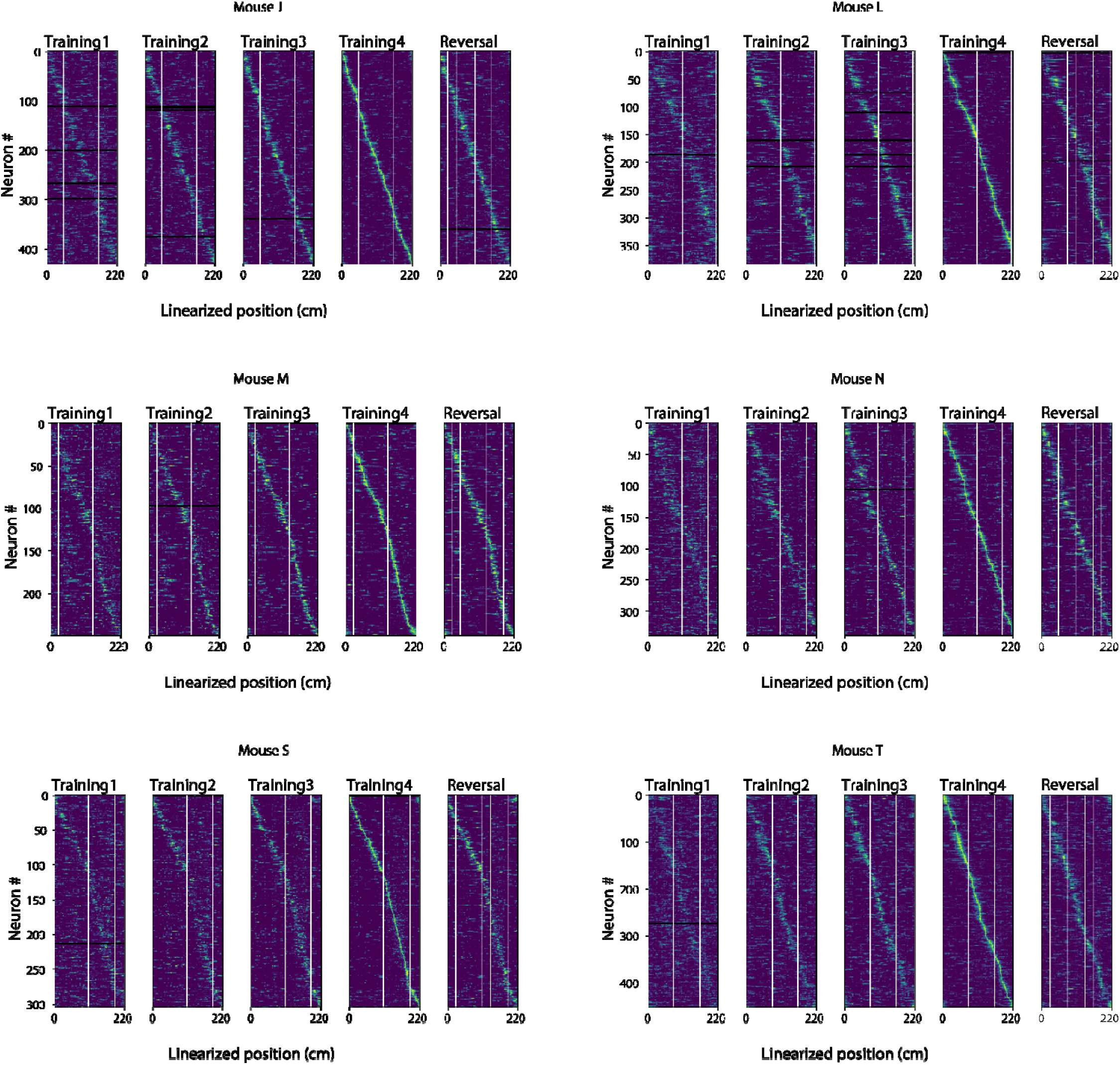
Spatial activity of neurons across days for all mice in Fig. 1. Trial-averaged activity of the same neurons recorded across Training and Reversal for 6 different mice. Neurons are sorted according to their field location on Training4. Only neurons that were registered across all five sessions are plotted. White rows correspond to neurons that had no activity during periods running above the speed threshold of 7 cm s_-1_. Yellow lines correspond to the rewarded port location. On the Reversal session, the faded yellow line corresponds to the reward ports on previous Training day.

**Supplemental Figure 4.**
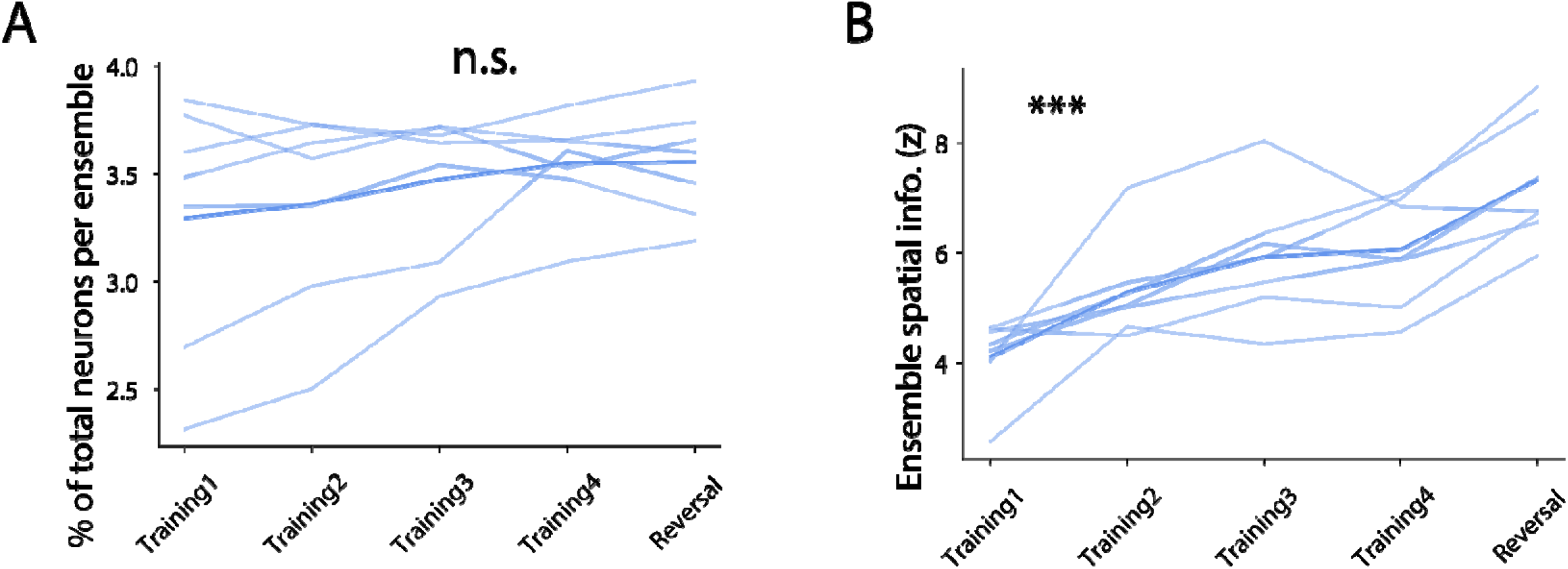
Ensemble properties in calcium imaging mice. A) Mean percentage of neurons in each ensemble. There was no significant difference across sessions (repeated-measures ANOVA F_4,24_=2.43, p=0.07). B) Average z-scored spatial information of ensembles for each mouse across sessions. Spatial information increased significantly over time (repeated-measures ANOVA F_4,24_=18.7, p<0.001).

**Supplemental Figure 5.**
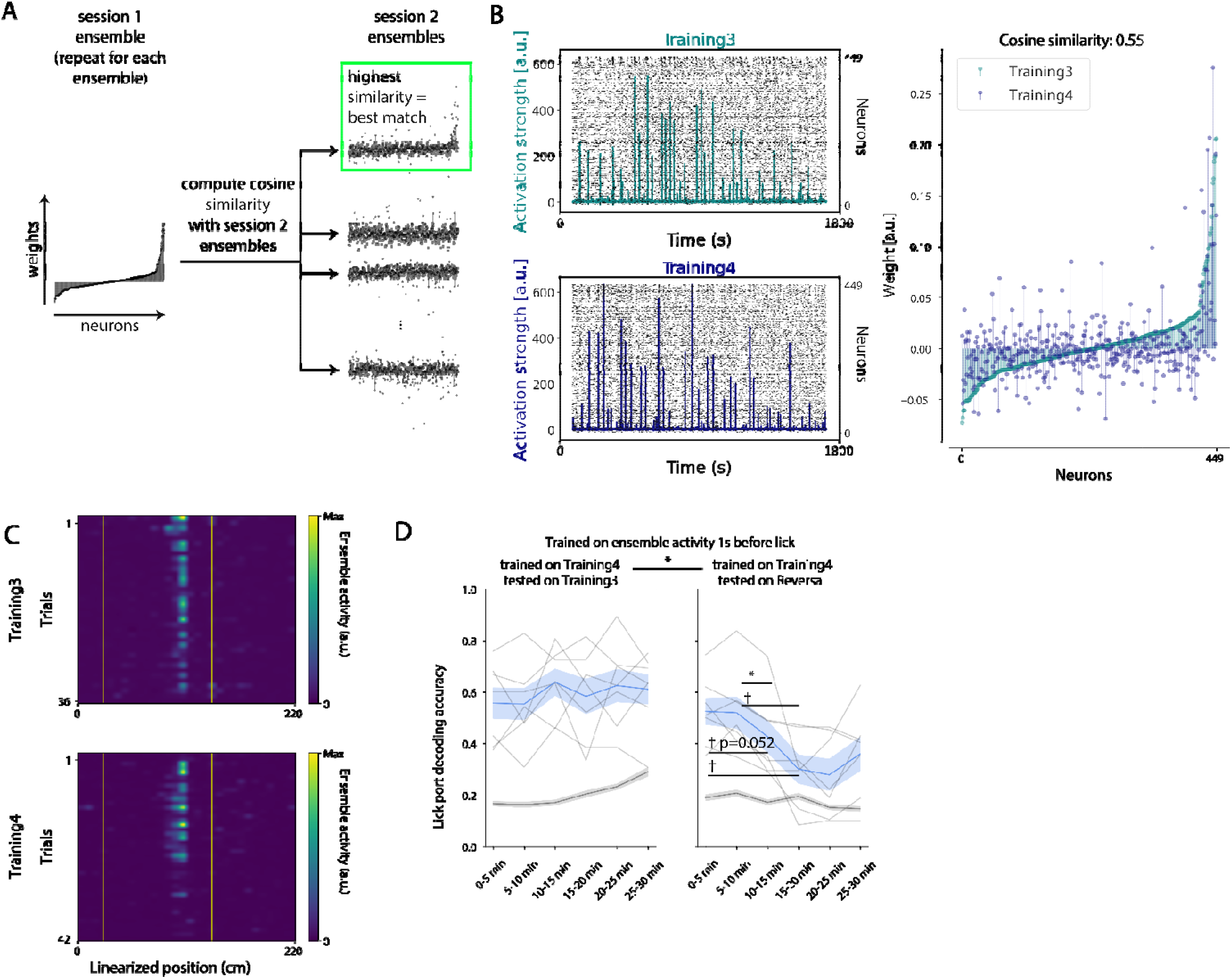
Cross-session decoding reveals a decaying ensemble code for lick port identity during memory-updating. A) Schematic illustrating our procedure for ensemble registration across sessions (see Methods). For each ensemble weight vector in session 1, we computed cosine similarities with every ensemble weight vector from session 2. The ensemble weight vector from session 2 with the highest cosine similarity (after exceeding a statistical threshold) gets registered to the session 1 ensemble. B) Example of a pair of ensembles that were registered across Training3 and Training4. Left, activation strengths of the two ensembles on their respective sessions. Right, weight vectors of the two ensembles, sorted by value on Training3. C) Activation of the same registered ensembles as B), plotted in space across trials. Yellow lines indicate rewarded port locations. D) Same as Fig 2g, but model training was performed on Training4 ensemble activity 1 s before the first lick of each trial on every reward port, then tested on either Training3 (left) or Reversal (right). Main effect of decoded session (repeated-measures ANOVA F_1,6_=7.78, p=0.032), but no main effect of time (F_5,30_=2.40, p=0.06). There was a significant interaction between decoded session and time (F_5,30_=4.69, p=0.0028). Post-hoc pairwise t-tests revealed significant decreases in decoding accuracy only on the Reversal session (*p<0.05 and †=0.052 after Sidak’s correction for multiple comparisons).

**Supplementary Figure 6.**
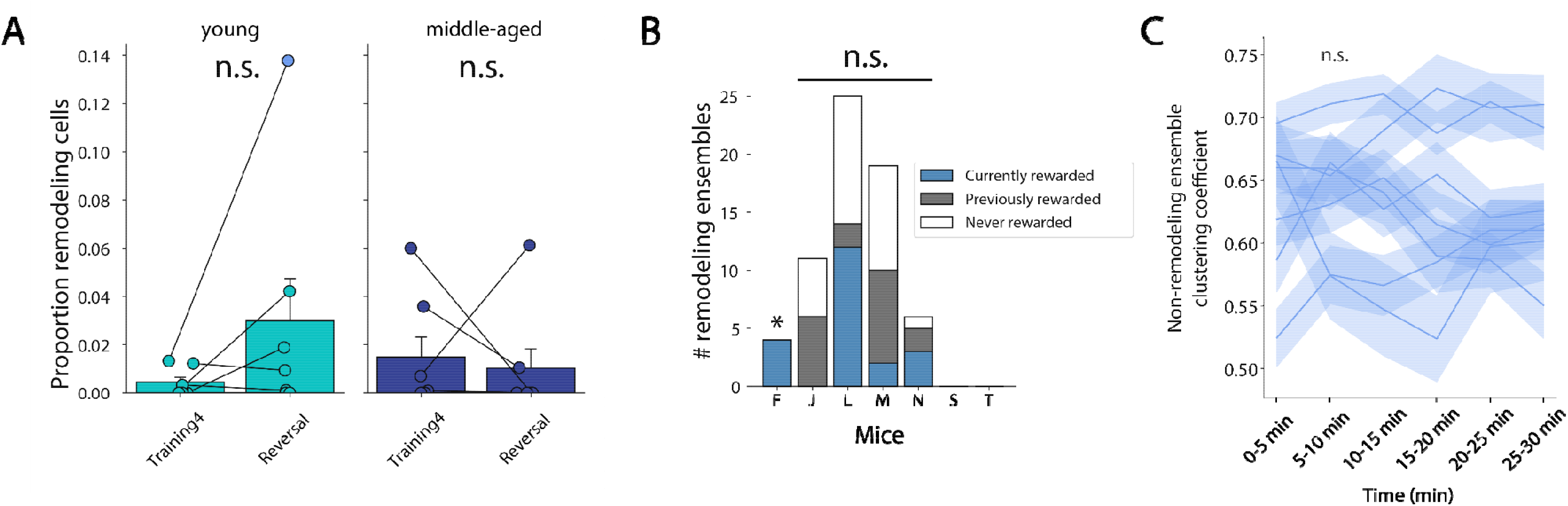
Properties of remodeling and non-remodeling ensembles and neurons. A) Proportions of remodeling neurons (neurons with decreasing activation strength out of all detected neurons) for Training4 and Reversal. The proportion of remodeling n urons is not statistically different during Reversal compared to Training4 in young mice (Wilcoxon signed-rank test, W=8.5, p=0.35) nor middle-aged mice (W=9.0, p=0.40). B) The proportion of remodeled ensembles located near currently rewarded, previously rewarded, and never rewarded ports during Reversal. Only one mouse (Mouse F) had a statistically uneven distribution of ensemble activity relative to reward ports (chi square test p<0.05 after Sidak correction for multiple comparisons). C) Clustering coefficients of non-remodeled ensembles across the Reversal session. There were no significant changes in 6 out of 7 mice (repeated-measures ANOVAs p>0.06), and no effect overall, averaged across all mice (F_5,30_=0.11, p=0.99).

**Supplemental Figure 7.**
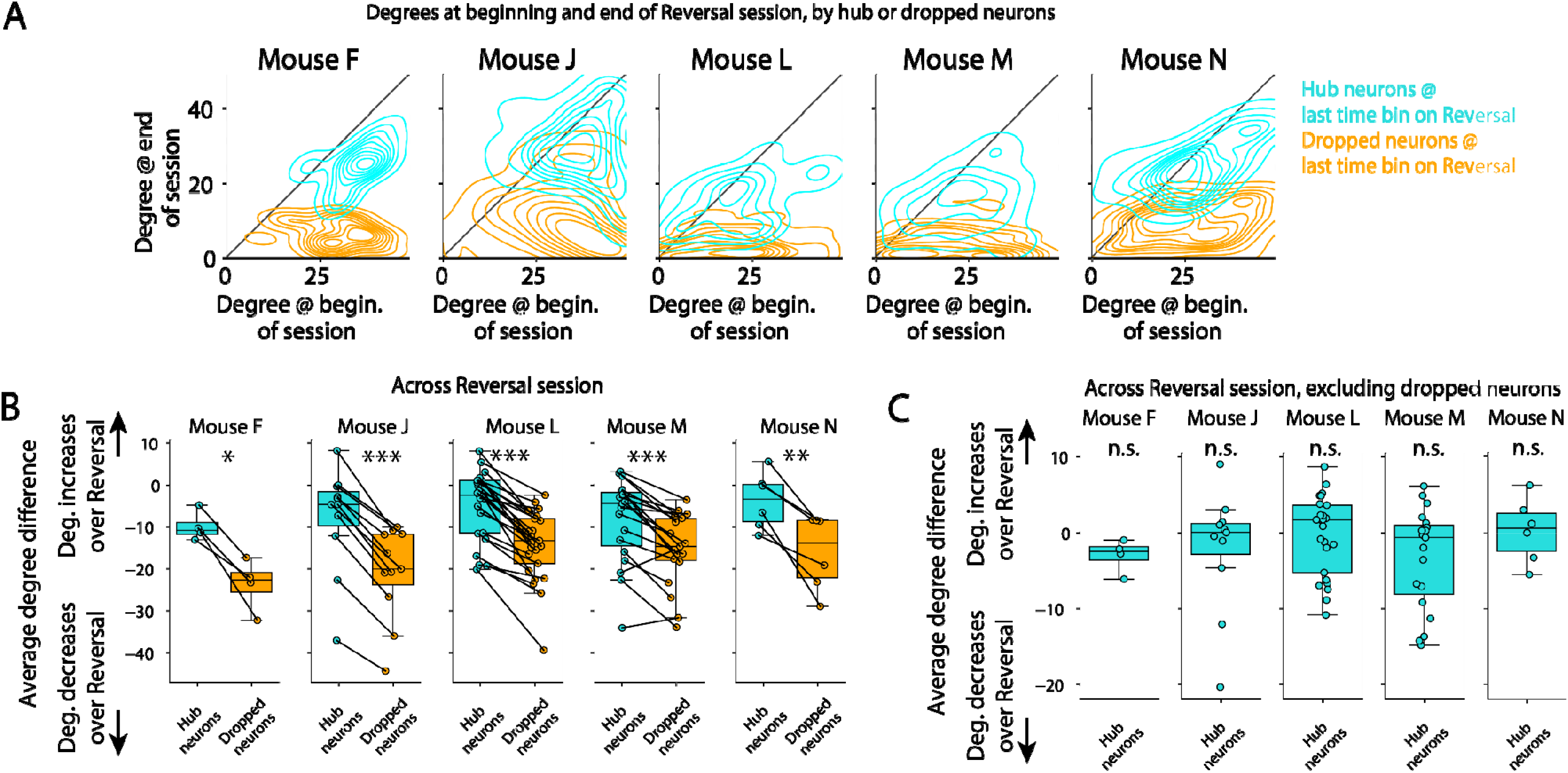
Co-activity differences between hub and dropped neurons. A) 2D histograms, as contour maps, of degree distributions at the beginning plotted against the end of the Reversal session, for both hub (cyan) and dropped (orange) neurons. Notice how the hub neurons stay along the diagonal, indicating stable degree across time, whereas the dropped neurons have lower degree at the end compared to the beginning of the session. B) Change in degree between the last time bin’s graph and the first time bin’s graph for each fading ensemble and mouse, split into hub and dropped neurons. In all mice, dropped neurons had higher decreases in degree across Reversal than hub neurons. (all p<0.05 after Benjamini-Hochberg corrections for multiple comparisons). C) Change in degree during Reversal, but when calculating the hub neurons’ degree difference, dropped neurons were excluded. In A), hub neurons significantly decreased their degree by the end of the Reversal session, but this was due to their disconnection from dropped neurons. If we exclude dropped neurons from the analysis, hub neurons did not decrease their degree over Reversal (one-sample t-tests compared to 0, all p>0.05 after Benjamini-Hochberg corrections for multiple comparisons), indicating that they remained co-active with the rest of the ensemble.

**Supplemental Figure 8.**
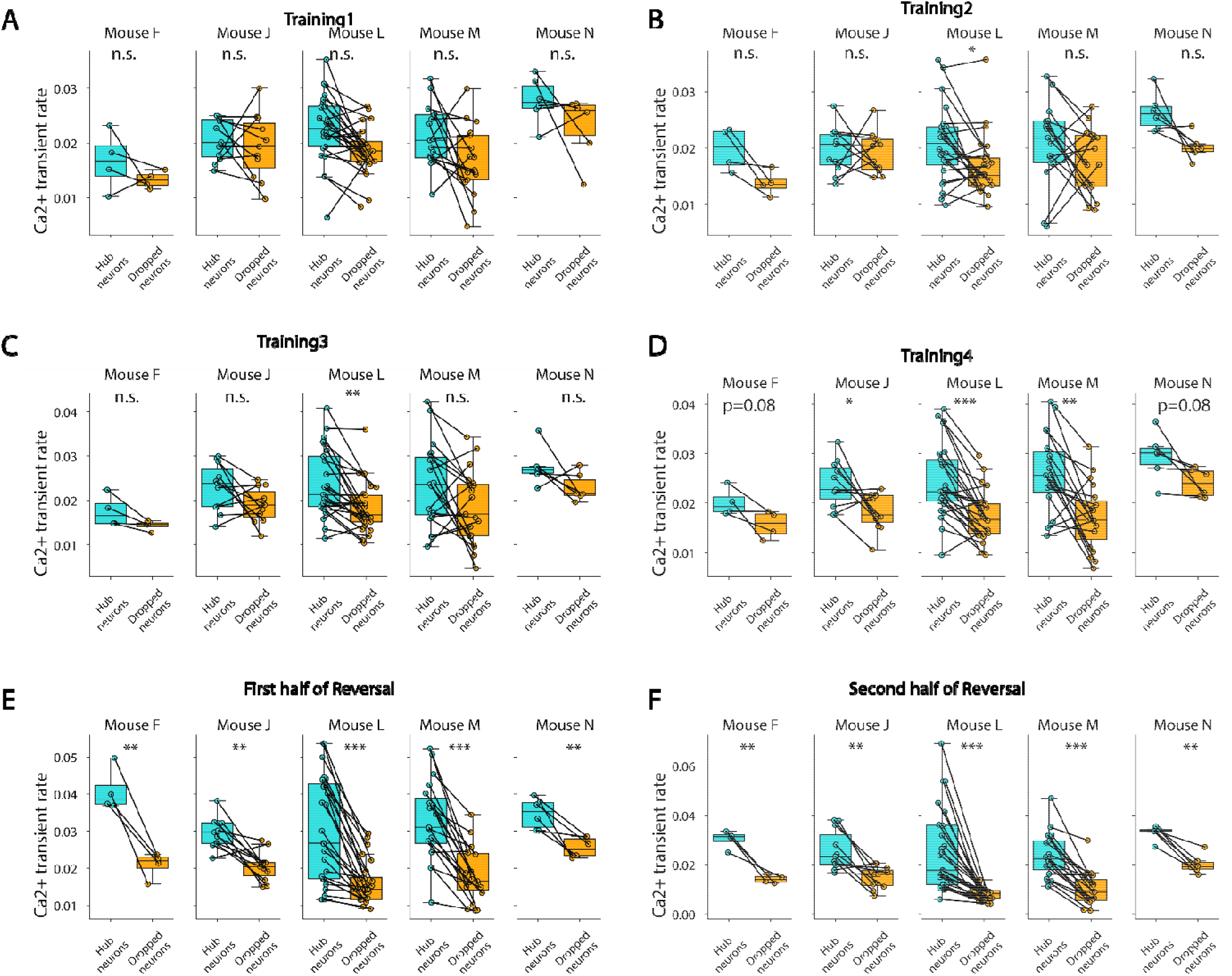
Activity rate differences between hub and dropped neurons. A) Average calcium transient rates of hub versus dropped neurons (classified during Reversal), on Training1. There were no differences in transient rates of hub versus dropped neurons in any mice (all p>0.05 after Benjamini-Hochberg corrections for multiple comparisons). B-F) Average calcium transient rates of hub versus dropped neurons (classified during Reversal). during Training2-Training4 and the first/second halves of the Reversal session. (*p<0.05, **p<0.01, ***p<0.001 after Benjamini-Hochberg corrections for multiple comparisons).

**Supplemental Figure 9.**
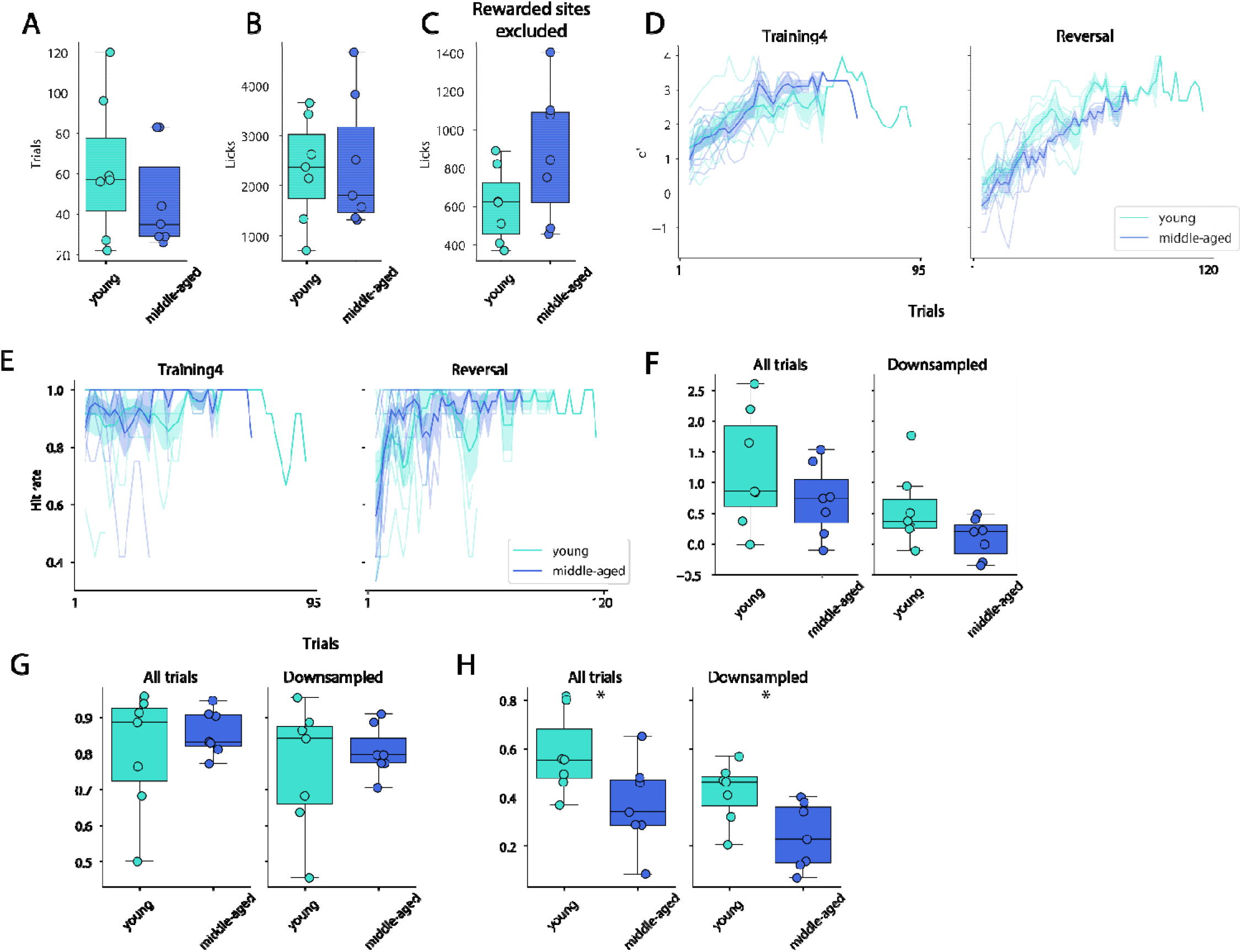
Behavioral differences in middle-aged mice. A) Trial counts in young and middle-aged mice during Reversal. There was no significant difference between the two groups (t_12_=0.94, p>0.05). B) Number of licks from young and middle-aged mice during Reversal. There was no significant difference between groups (t_12_=-0.18, p>0.05). C) Number of licks during Reversal, excluding rewarded ports. There was no significant difference between groups (t_12_=-1.78, p>0.05). D) Behavioral d’ of young and middle-aged mice during Training4 and Reversal. There was a significant difference between young and middle-aged mice during Reversal (two-way ANOVA F_1,57_=35.6, p<0.001), but not Training4 (F_1,45_=0.02, p=0.9). E) Hit rate during Training4 and Reversal. There were no significant differences between young and middle-aged mice in both Training4 (two-way ANOVA F_1,45_=1.23, p=0.27) and Reversal (F_1,57_=3.15, p=0.07). F) Behavioral d’ for young and middle-aged mice averaged across all trials (left) and after downsampling trials to match trial counts across the two groups (right). There were no significant differences across the two groups (t-tests, p>0.05). G) Hit rate of young and middle-aged mice averaged across all trials, and after downsampling. There were no significant differences across the two groups (t-tests, p>0.05). H) Correct rejection rate of young and middle-aged mice averaged across all trials, and after downsampling. There was a significant difference between young and middle-aged mice when considering all trials (left, t_12_=2.24, p=0.045) and downsampled trials (right, t_12_=2.61, p=0.023).

